# Coenzyme A is bound to tafazzin – a paradigm change for transacylation

**DOI:** 10.64898/2026.02.05.703992

**Authors:** José Guadalupe Rosas Jiménez, Jonathan Schiller, Janet Vonck, Gerhard Hummer, Volker Zickermann

## Abstract

Cardiolipin (CL) is the signature phospholipid of mitochondria. In an obligatory remodeling process, the mitochondrial transacylase tafazzin exchanges its acyl chains to create the highly unsaturated, mature form of CL. Tafazzin dysfunction causes Barth syndrome, a severe multisystem disorder. We determined the structure of tafazzin at a resolution of 2.2 Å using cryo-electron microscopy (cryo-EM). Until now, the tafazzin reaction has been thought to be independent of coenzyme A (CoA). However, our structure clearly shows an acyl-CoA molecule bound to tafazzin. To decipher how substrates bind to the active site, we combine cryo-EM with structure predictions and molecular dynamics simulations, giving us detailed insights into a transacylation mechanism mediated by CoA. By providing molecular explanations of tafazzin dysfunction caused by pathogenic mutations, we gain a molecular understanding of Barth syndrome.

## Introduction

Mitochondria have a characteristic double membrane structure. The inner membrane is highly folded into cristae, which are the site of ATP generation by oxidative phosphorylation (OXPHOS). Cardiolipin (CL) accounts for up to 15% of the phospholipids in the inner mitochondrial membrane and is essential for its distinctive structure and the functionality of the OXPHOS complexes (*1-5*). CL has a unique architecture with two phosphatidic acid units connected by a glycerol molecule. The lipid is synthesized in the inner mitochondrial membrane, followed by an obligatory remodeling process in which the composition of its four acyl chains is altered (*2, 6*). Typically, this results in cardiolipin with a higher content of unsaturated fatty acids and thus altered physical properties.

Cardiolipin remodeling is catalyzed by tafazzin, an enzyme with a broad substrate spectrum that transfers fatty acids from phospholipids to lyso-phospholipids, e.g., to monolysocardiolipin (MLCL) (*7*). Together with a selection of lyso-phospholipid acyltransferases (LPLATs) and glycerol-3-phosphate acyltransferases (GPATs), tafazzin belongs to the family of 1-acylglycerol-3-phosphate O-acyltransferases (AGPATs) that is characterized by a HisX_4_Asp active site motif (*8, 9*). The canonical acyltransferases of the AGPAT family transfer an activated acyl chain appended to coenzyme A (CoA) or an acyl carrier protein (ACP) to glycerol-3-phosphate or a lyso-phospholipid compound. By contrast, the reaction of tafazzin converts a phospholipid-lysophospholipid pair into a corresponding pair with an altered acyl chain composition. Biochemical experiments demonstrated that this transacylase reaction catalyzed by tafazzin does not involve CoA or an ACP as a donor or acceptor of the transferred acyl chain (*7, 10*), leaving it unclear how the chain transfer is activated. Also, the substrate specificity of tafazzin is discussed controversially. *In vivo*, tafazzin is responsible for the formation of CL with unsaturated acyl chains, and a preference for the transfer of unsaturated fatty acids has been observed in transacylation assays *in vitro* (*7, 10, 11*). However, it has also been shown that the reaction of tafazzin is context-dependent, leading to the conclusion that CL rearrangement is a self-organizing process that is not determined by the substrate specificity of tafazzin, but rather by the thermodynamic driving force at the site of tafazzin action (*12*). This mechanism is thought to ensure that, depending on local conditions, the lipid tail composition with the lowest free energy is produced.

In humans, dysfunction of tafazzin causes Barth syndrome, a rare inherited multisystem disorder (*13-15*). The characteristic symptoms are cardiomyopathy, skeletal myopathy, neutropenia, and growth retardation (*16, 17*). An increased MLCL/CL ratio is the most important diagnostic marker and CL with abnormal acyl chain composition is typically found (*18*). Various animal model systems to study the disease have been established, but disease-specific treatment options are still under development (*19*). The impact of different classes of pathogenic missense mutations on enzyme stability and activity has been evaluated in a baker’s yeast model (*20*). However, the molecular mechanism of tafazzin has remained elusive due to the lack of structural information. We have recently determined the 3.2 Å resolution cryo-EM structure of tafazzin as a component of an assembly intermediate of respiratory complex I from the aerobic yeast *Y. lipolytica* (*21*). We could show how tafazzin associates with the matrix side of the inner mitochondrial membrane and we provisionally assigned the function of several conserved residues in catalysis and lipid binding. Here, we present the structure of tafazzin at improved resolution and show that, unexpectedly, an acyl-CoA molecule is bound to tafazzin as isolated. Our new structural data combined with structure predictions and molecular simulations lead us to a detailed mechanism of tafazzin-mediated transacylation in which CoA plays a central role.

## Results

### High-resolution structure of tafazzin

We have previously shown that in the aerobic yeast *Y. lipolytica*, tafazzin is a component of the early assembly intermediate of the P_P_ module of respiratory complex I (*21, 22*). We were now able to determine the structure of a yet uncharacterized P_P_ module assembly intermediate that comprises seven complex I subunits, assembly factors NDUFAF1, CIA84, TPR50, CMC1L, and tafazzin. Improvements in sample preparation, data collection and data processing resulted in an excellent cryo-EM map with a resolution up to 2.2 Å (fig. S1). The structure of the assembly intermediate (fig. S1, movie 1) will be described elsewhere, and we focus here exclusively on tafazzin.

*Y. lipolytica* tafazzin has a mass of 43 kDa and consists of 372 residues. Our model of tafazzin comprises 358 residues and 159 water molecules; three lipid molecules sit at the membrane/tafazzin interface. The overall structure of tafazzin (Fig. 1A) and its interactions with assembly factors CIA84 and NDUFAF1 are very similar to our previously reported model (*21*). Briefly, the core of tafazzin is a seven-stranded β-sheet that is surrounded by α-helices. Helix α-2 is a long amphipathic helix that lies on the membrane surface. The active site residues of the HisX_4_Asp motif are in the loop connecting helix α-6 and β-strand 3. As in our previous structure, the sequence stretch between position 106 and 119 showed only weak density in the map and was omitted from the structure. An AlphaFold model (*23*) of tafazzin from *Y. lipolytica* suggested an arrangement of two short helices at this position that are approximately perpendicular to each other (*21*). One of them points downwards to the membrane surface and we speculate that this structural element is disturbed by the detergent micelle. In agreement with our previous study, we found a conspicuous density running from the surface of the protein about 35 Å above the membrane to a position close to the active site His76, which we had previously interpreted as an alternative protein fold of the disordered 106 to 119 sequence stretch (*21*). Thanks to the high quality of the new map (fig. S2) it was now straightforward to unequivocally model a CoA molecule into the density with its reactive thiol group close to the active site. A further inspection of the map showed that from here a weaker density continued towards membrane associated helix α-2. We interpret this density as an acyl chain appended to the CoA molecule; however, the weaker density indicates a higher mobility and/or lower abundance of the acyl chain (fig. S2). Please note that we did not add CoA or CoA derivatives before or during protein purification or sample preparation, so we did not alter the native state of tafazzin with regard to CoA binding. Intriguingly, the map has additional density features in the active site and at the protein membrane interface, which may originate from a bound substrate lipid molecule; but they are too weak for reliable model building.

**Figure 1.**
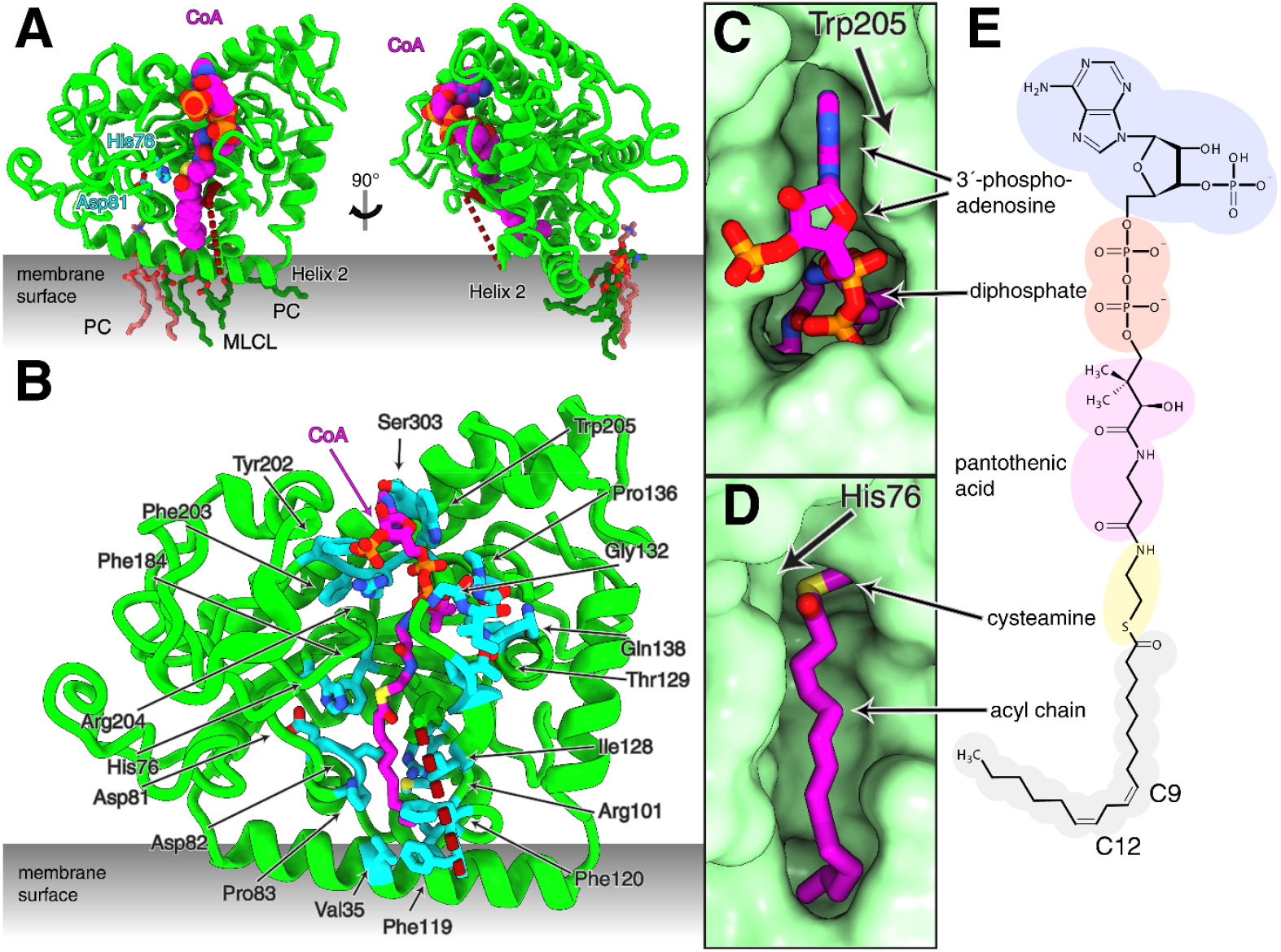
Cryo-EM structure of *Y. lipolytica* tafazzin at 2.2 Å resolution. (A) Two views of tafazzin with acyl-CoA shown in sphere representation; unmodeled sequence stretch between position 106 to 119 indicated by red dashed line. Tafazzin is part of an assembly intermediate of respiratory complex I not shown here for clarity (see figs. S1 and S8 and movie 1). (B) Residues involved in coordination of acyl-CoA in tafazzin. For a more comprehensive presentation see movie 1. (C,D) Surface representation of the acyl-CoA binding pocket. (C) CoA binding pocket viewed from the matrix side. (D) Position of the thioester bond and acyl chain in the acyl chain binding pocket, cryo-EM density allowed modeling of a C14 chain. (E) Lewis structure of linoleoyl-CoA.

### Binding of acyl-coenzyme A to tafazzin

Coenzyme A can be dissected into four parts (Fig. 1E): a 3’-phosphoadenosine head group connected by a diphosphate to pantothenic acid (vitamin B5), followed by cysteamine. The SH group of the cysteamine moiety can bind an acyl group in a thioester linkage. In our cryo-EM structure of tafazzin, the adenine ring of the CoA head group is embedded in a tight binding pocket with a variety of interactions (Fig. 1B, C and movie 1). Most notable is a stacking interaction with the indole ring of the highly conserved Trp205 (fig. S3 A,B and fig. S4). The amino group and nitrogen atoms of the purine ring at positions 1 and 7 interact with the protein backbone and with the side chain of Ser303. The ribose unit is located on the protein surface. The phosphate in the 3’ position and the α-phosphate of the diphosphate unit interact with Arg204. A basic residue at this position, usually lysine, is highly conserved in tafazzin (fig. S4). The diphosphate also forms hydrogen bonds with the indole nitrogen of Trp205 and the side chain of Tyr202, as well as with backbone amide nitrogens of a glycine-rich sequence extending from positions 132 to 136. Strictly conserved Gln138 points towards the β phosphate and stabilizes the glycine-rich loop by a backbone interaction with Thr129. Pantothenic acid engages in several polar and nonpolar interactions with the protein. The methyl groups point towards the side chains of Ile128 and Met141, respectively. The hydroxyl group forms an intramolecular hydrogen bond with the diphosphate, while the other polar groups mainly form backbone interactions. Several buried water molecules mediate contact between CoA and the protein structure. Strikingly, the thioester bond between the cysteamine moiety and the appended acyl chain is close to His76 that together with Asp81 forms the strictly conserved HisX_4_Asp motif defining the active site of the AGPAT family (Fig. 1B,D and movie 1). The well-resolved 14 carbon atoms of the acyl chain connected to the cysteamine group insert into a binding pocket that is predominantly lined with hydrophobic residues (Fig. 1B,D and fig. S3C). A ∼10-Å segment formed by carbon atoms 1-9 extends straight from the active site in the direction of helix α-2; then, the chain turns sharply towards the interior of the protein (Fig. 1B,D and fig. S3C and movie 1). The first four carbon atoms of the acyl chain are located near the highly conserved Phe184, which has a close stacking interaction with the imidazole ring of the active site His76. The pocket continues with Ala103 and Val125, followed by Phe119 and the highly conserved Phe120. Together with Val35 from the α-2 helix, these two phenylalanines form the outer boundary of the sharp bend of the acyl chain, while highly conserved Pro83 and Gly123 together with Ile32 build the entrance to the terminal part of the pocket that is buried deep in the protein. Met100 and neighboring residues constitute the end of the pocket. Surprisingly, the ion pair formed by the highly conserved residues Asp82 and Arg101 is also part of the otherwise hydrophobic binding pocket (fig. S3C). Asp82 is followed by the fully conserved Pro83, forming a *cis* peptide bond that brings the Asp side chain in position to form a tight salt bridge with Arg101.

Overall, our cryo-EM structure clearly shows that a CoA molecule is bound to tafazzin from *Y. lipolytica*. Although we did not add CoA during purification or sample preparation, the cryo-EM density for CoA is strong, indicating that occupancy of the binding site is high. The conservation of important residues strongly suggests that CoA binding is a general feature of tafazzin (see also below). We find a binding pocket for an acyl chain attached to CoA, but weaker density complicates modeling of the full-length acyl chain. The map does not allow the modeling of complete (lyso)-lipid molecules in or close to the active site. Therefore, we extended our approach and used AlphaFold3 structure predictions and molecular dynamics (MD) simulations to further investigate the binding of CoA and substrate molecules for the transacylation reaction.

### AlphaFold 3 structure predictions support binding of acyl-CoA and MLCL

AlphaFold 3 predicts the structure of tafazzin from *Y. lipolytica* in very good agreement with the cryo-EM data (Fig. 2A). The excellent match of the binding sites for CoA is particularly noteworthy, as the training dataset did not contain any member of the AGPAT family with bound coenzyme (*24*). The 18:2 acyl chain of the linoleoyl-CoA included in the AlphaFold 3 model extends beyond the 14 C atoms we could confidently model in the cryo-EM density. Notably, the last ∼8 C atoms were predicted with high confidence inside the hydrophobic cavity that we had already observed in the cryo-EM structure. A bend of the linoleoyl chain at the position of the Δ-9 unsaturation is clearly visible (Fig. 2B,C) and follows a curve defined by the conserved residues Phe119 and Phe120. Interestingly, the guanidinium group of the Asp82/Arg101 salt bridge is close to the double bonds of the linoleic acid chain. This remarkable structure requires the highly conserved Pro83 to be in the unusual *cis* configuration. At the bottom of the cavity Trp86 and Met100 engage in hydrophobic contacts with the last carbon atom of the linoleoyl chain.

**Figure 2.**
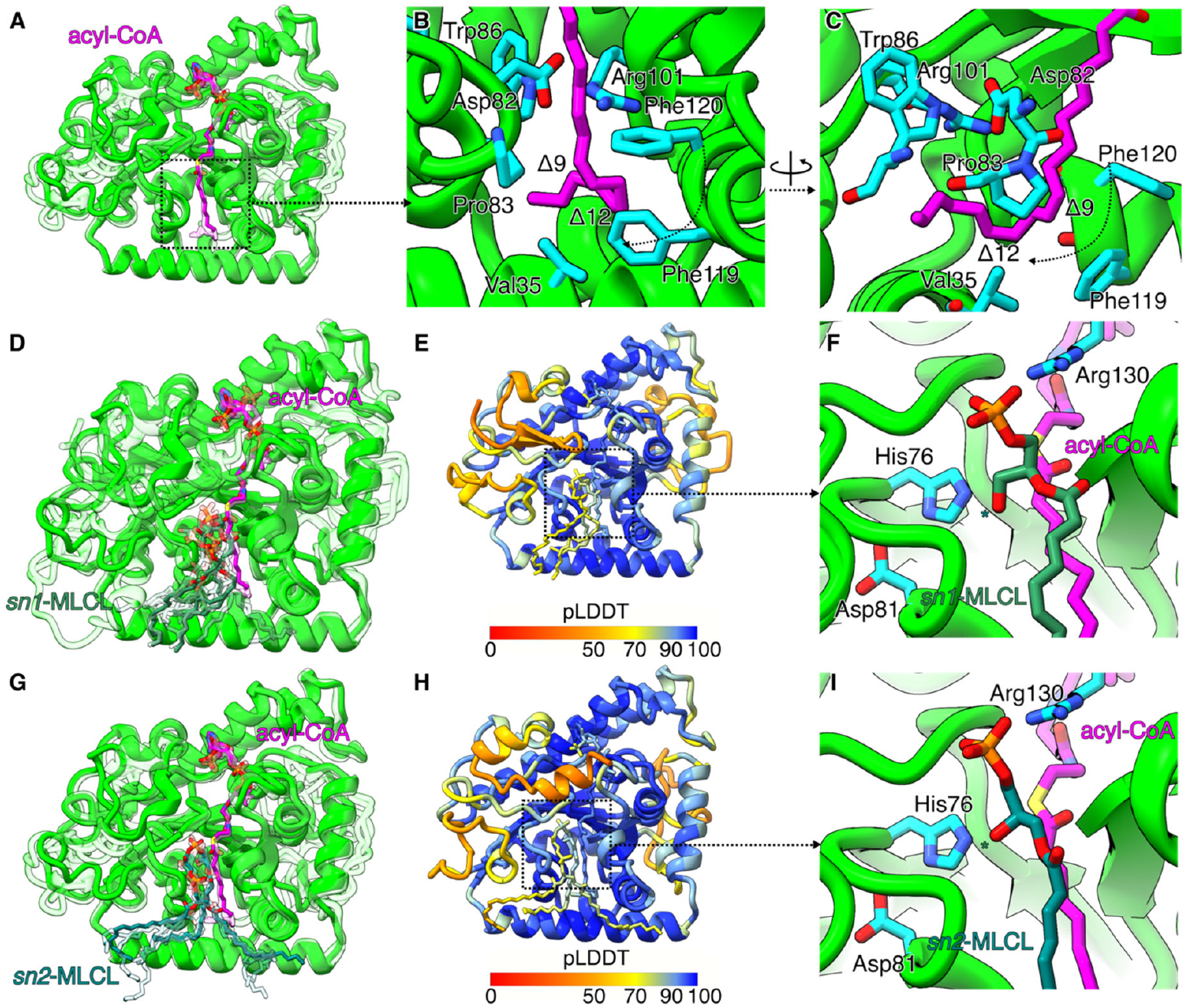
AlphaFold 3 structure prediction for tafazzin in complex with acyl-CoA and monolysocardiolipin. (A) Superposition of the five structures with the highest ipTM score for acyl-CoA (transparent), aligned with respect to the experimental structure (solid). (B,C) Detailed view of the hydrophobic tunnel where the acyl chain attached to CoA is predicted to bind; note the prominent kink of the acyl chain close to Phe119 and Phe120. (D,E) Prediction of how *sn*1-hydroxy-MLCL binds to the tafazzin-acyl-CoA complex. The solid structure of MLCL is the prediction with highest ipTM score. Atoms are colored according to the local pLDDT values. (F) Detailed view of the catalytic site when *sn*1-hydroxy-MLCL is bound to tafazzin. Only the PA1 subunit of MLCL is shown for clarity. The star indicates the hydroxyl group involved in the acyl transference reaction. (G,H) Prediction of how *sn*2-hydroxy-MLCL binds to the tafazzin-acyl-CoA complex. The solid structure of MLCL is the prediction with highest ipTM score. Atoms are colored according to the local pLDDT values. (I) Detailed view of the catalytic site when *sn*2-hydroxy-MLCL is bound to tafazzin. Only the PA1 subunit of MLCL is shown for clarity. The star indicates the hydroxyl group involved in the acyl transfer reaction.

To unravel the binding site of the acceptor lysolipid, we included MLCL with three palmitoyl chains in the AlphaFold 3 modeling and the reactive hydroxyl group either in the *sn-1* (Fig. 2D-F) or *sn-2* position (Fig. 2G-I). For the description of MLCL, we introduce the nomenclature illustrated in supplemental figure S7. Briefly, the lysophosphatidic acid subunit is denoted as PA1, and the complete phosphatidic acid subunit is denoted as PA2. The pLDDT scores indicate a high confidence in the predicted conformation of the PA1 subunit, independently of the position of the reactive hydroxyl (Fig. 2E,H). This moiety is in contact with the catalytic site and close to linoleoyl-CoA. By contrast, the acyl chains of the PA2 subunit show a lower confidence score, most likely because the membrane is not taken into account in AlphaFold 3 modelling. Remarkably, in all 100 generated predictions, the MLCL head group is bound to the expected catalytic site. The reactive hydroxyl group in MLCL is consistently placed within a distance of ∼2.90 Å and ∼4.28 Å to the ε-nitrogen atom of His76 and the thioester carbon in linoleoyl-CoA, respectively. Strikingly, AlphaFold predicts this short distance between reactive groups for both positions of the lysophospholipid hydroxyl acceptor (Fig. 2F,I). This suggest that the flexibility of the glycerol backbone is high enough to adapt to the catalytic site depending on the position of the hydroxyl group. Therefore, AlphaFold 3 predictions show all reactants in an optimal position for acyl chain transfer, but it is not yet clear how the negatively charged transition state is stabilized.

### MD simulations reveal how MLCL is recruited from the membrane

We selected the best scored AlphaFold prediction as starting structure for MD simulations of the protein in a membrane/solvent environment. In our simulations, linoleoyl-CoA stayed tightly bound to the protein. The adenine ring and the pantothenic acid moiety of CoA showed particularly small positional fluctuations (fig. S5A-C). The time series of the root mean squared deviation (RMSD) with respect to the initial frame form a plateau during a wide range of the simulation time (fig. S5B-D), indicating minimal changes in the conformation (fig. S5A). The complex is stabilized by the prominent stacking interaction between the adenine ring of the CoA head group and Trp205, and by an ionic interaction between Arg204 and the 5’ and 3’ phosphates of ribose (fig. S5E). Additionally, persistent hydrogen bonds between the adenine ring and the backbone oxygen of Phe203 and the side chain hydroxyl of Ser303 were observed during the simulations. These results show that this site is complementary and specific to the nucleotide moiety of CoA. A detailed trajectory sample of our simulations, highlighting the binding of acyl-CoA, is presented in Movie S2.

Remarkably, also the less constrained acyl-chain conformation of linoleoyl-CoA in the AlphaFold 3 model is preserved in the MD simulations (fig. S5A). Atoms C1-9 of the linoleoyl chain extend from the catalytic site towards the membrane and, at the Δ-9 double bond, the acyl chain bends towards the hydrophobic cavity that binds atoms C10-18 (fig. S6). Histograms of the center-to-center distance of Δ-9 and Δ-12 double bonds to Phe119 and Phe120 exhibit peaks at ∼5 Å, suggesting the formation of π-π interactions between the double bonds in the acyl chain and the aromatic residues (fig. S6 B,C). Particularly, the unsaturation at position Δ-9 shows interactions with both aromatic residues. The presence of the Δ-9 double bond has been highlighted previously as an important factor of acyl chain recognition (*10, 11*). Furthermore, the conserved Pro83 contributes to acyl chain bending by forming persistent contacts at the position of the double bonds, indicating a role in the selection of unsaturated acyl chains at the level of acyl-CoA binding (fig. S6D). *In vitro* experiments with *trans*-unsaturated acyl chains showed a decreased transacylation activity (*11*), demonstrating the importance of the double bond stereochemistry for acyl chain binding, in line with our simulation results.

The MD simulations also shed some light on chain-length preference. Biochemical experiments have demonstrated that acyl chains longer than 18 carbon atoms are poorly transferred (*7*). In our MD simulations, a C-18 acyl chain reaches all the way to the bottom of the cavity, contacting residues Trp86 and Met100, indicating that binding of longer acyl chains would cause steric clashes (fig. S6 E,F). Indeed, the minimum distance between the conserved residue Trp86 buried deep into the cavity and the linoleoyl chain exhibits an average of ∼3 Å in simulations, suggesting that longer chains may be less easily inserted into the cavity.

The MD simulations also reveal the factors governing MLCL recruitment to tafazzin (Mov S3). AlphaFold 3 predicted the conformation of the PA1 subunit with higher confidence than the conformation of PA2. Our MD simulations show that acyl chains in the PA2 subunit are indeed more flexible than the single acyl chain of the PA1 subunit. Lipid binding concentrates in three distinct regions: A1, where the single acyl chain of PA1 binds; A2, which binds to the acyl chains in the PA2 subunit; and G, where the phosphatidylglycerol head group binds (Fig. 3A,B). The A1 site is lined by Val79, Leu80, Val84, Pro83, Phe42, Cys39, Met38, Val35, and Leu34 (Fig. 3C). The A1 hydrophobic cleft extends from the catalytic site, beginning at Val79, all the way to the protein-membrane interface at helix 2, thereby providing a passageway for lipid extraction from the membrane into the catalytic site approximately 10 Å above the membrane. Indeed, an approximately 8 Å long density feature most likely originating from a bound substrate molecule is seen in the cryo-EM map close to the side chains of residues Leu80, Val84, Phe42, Met38, and Val35. The A1 cleft is also adjacent to the binding site of the acyl-chain of acyl-CoA, inducing the formation of extensive contacts between the acyl chains in the PA1 subunit of MLCL and acyl-CoA. These close contact interactions favour the proximity between the chemical groups that participate in the reaction.

**Figure 3.**
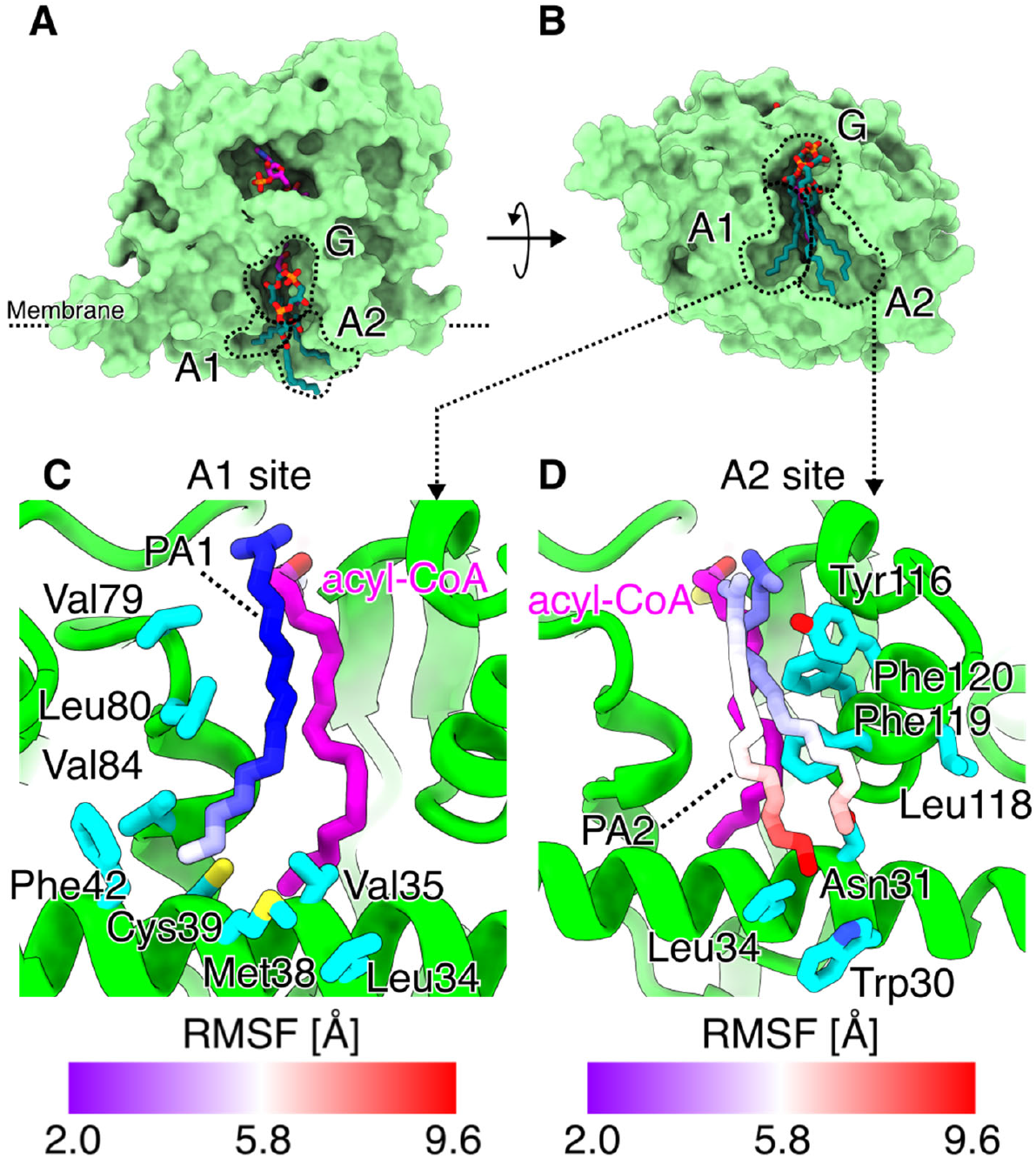
Monolysocardiolipin binding sites in MD simulations. (A,B) Side (A) and bottom view (B) of the MLCL binding site in MD simulations. Binding regions A1, A2, and G are indicated. (C) Structure and residues forming the A1 site, which recognizes the acyl chain in the PA1 subunit of MLCL. (D) Structure of the A2 site. Both acyl chains in the PA2 subunit of MLCL can form contacts with this site. MLCL acyl atoms are colored by RMSF values in (C,D).

The A2 site presents a comparably shallow hydrophobic patch for lipid binding, formed by residues Leu34, Trp30, Asn31, Tyr116, Leu118, Phe119, and Phe120 (Fig. 3D). This corresponds in part to the flexible sequence stretch close to the membrane surface (residues 106 – 119) that is not resolved in the cryo-EM structure. Acyl chains in the PA2 subunit of MLCL form transient but frequent interactions with this site. Interestingly, AlphaFold 3 predicts that the *sn*-1 acyl chain is bound in the A2 region. However, in the MD simulations we have observed conformations in which either one or the other PA2 acyl chain can bind in the A2 cleft.

Finally, the MLCL headgroup is bound by the more hydrophilic G region formed by residues Ile109, Asp108, Ala106, Arg130, Tyr188, Val189, and His76. Notably, this binding site recognizes the glycerol head group of MLCL by forming electrostatic interactions between the highly conserved Arg130 and the phosphate group in the PA1 subunit of MLCL (Fig. 4). Besides lipid binding, this region also encloses the reactive thioester group of acyl-CoA and the catalytic dyad His76/Asp81, forming an appropriate environment for the acyl transfer reaction.

**Figure 4.**
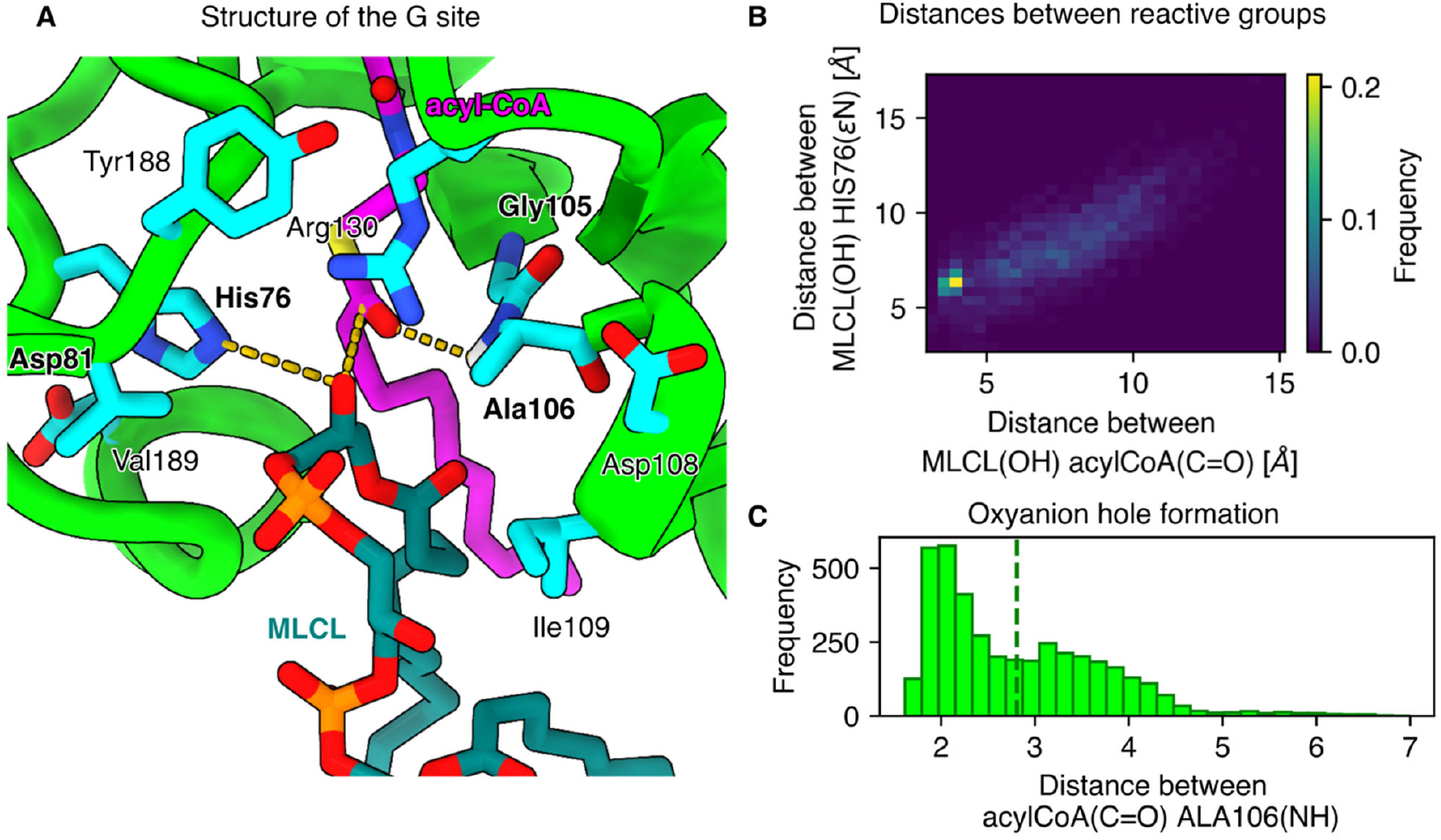
Monolysocardiolipin accesses the catalytic site in MD simulations. (A) Structure of the catalytic G site observed in MD simulation. All elements required by the reaction are contained in this site: His76 acting as an activator, the thioester group of acyl-CoA ready for a nucleophilic attack, and the hydroxyl group of MLCL acting as a nucleophile. (B) Two-dimensional histogram of the distances between the reactive hydroxyl group and His76 and between the thioester carbon atom in acyl-CoA. The high flexibility of the glycerol headgroup allows adaptation of the lipid to the catalytic site, according to the position of the free hydroxyl group. (C) Histogram of the distances between the oxygen atom in the thioester group of acyl-CoA and the amide backbone between residues Gly105 and Ala106, forming a geometry consistent with an oxyanion hole.

It is noteworthy that the configuration of substrates in the active site meets all requirements for a productive enzyme-substrate complex (Fig. 4A), similar to what has been shown or proposed previously for other members of the AGPAT family (*25, 26*). In the MD simulation, the hydroxyl in unit PA1 of MLCL is simultaneously close to residue His76 and the thioester carbon in acyl-CoA, at distances of ∼6.3 and ∼3.9 Å, respectively (Fig. 4B). Thermal fluctuations bring these groups into contact in the presumed active site region. The distribution of the two distances is positively correlated, indicating that when the MLCL hydroxyl group approaches the thioester carbon, His76 also gets closer to the reactive groups. Furthermore, the flexibility of the PA1 glycerol headgroup observed in simulations indicates that the free hydroxyl group can rotate and adapt to the catalytic site, independently if it is at the *sn1* or *sn2* position. The positive charge of the protonated His76 is balanced by Asp81, the other canonical active site residue. Interestingly, the carbonyl oxygen of the CoA thioester points towards the backbone of residues Gly105 and Ala106 (Fig. 4C). This arrangement is typical for an oxyanion hole that stabilizes a transition state with a negative charge on an oxygen atom (*27, 28*) .

### AlphaFold 3 prediction and MD simulations of donor lipid binding

The prediction of a plausible enzyme substrate complex between MLCL and linoleoyl-CoA by AlphaFold 3 is striking. Given that tafazzin is a transacylase, a corresponding complex should form between a phospholipid acyl chain donor and tafazzin. Since CoA appears to be an integral component of the enzyme, we pursued structure prediction for a phosphatidylcholine (PC) molecule bound to tafazzin with CoA in its binding pocket (Fig. 5). The CoA molecule is predicted essentially in the same conformation as the CoA molecule carrying an acyl group. Strikingly, the acyl chain at the *sn*-2 position of PC is docked in the same cavity where the tail of acyl-CoA binds (Fig. 5A). By contrast, the acyl group at the *sn*-1 position is predicted to bind in a conformation equivalent to the acyl chain in the PA1 subunit of MLCL (Fig. 5C,E). Results from MD simulations, however, show that the *sn*-1 acyl group of PC is constantly exchanged between the A1 and A2 binding sites (Mov S4). This dynamic suggests that A1 and A2 sites are hotspots for interactions with acyl chains in the membrane. Similar to MLCL, the G site prevents the lipid from escaping to the membrane by interacting with the phosphatidic acid backbone of PC (Fig. 5D). This includes strong electrostatic interactions of the phosphate group of PC with the strictly conserved Arg130. Notably, the acyl chain at the *sn*-2 position has the lowest RMSF values (Fig 5E), in comparison with the highly flexible chain at *sn*-1. Therefore, PC is tightly bound to the protein by occupying the same cavity where the acyl chain attached to CoA would bind in the complex with the lysolipid.

**Figure 5.**
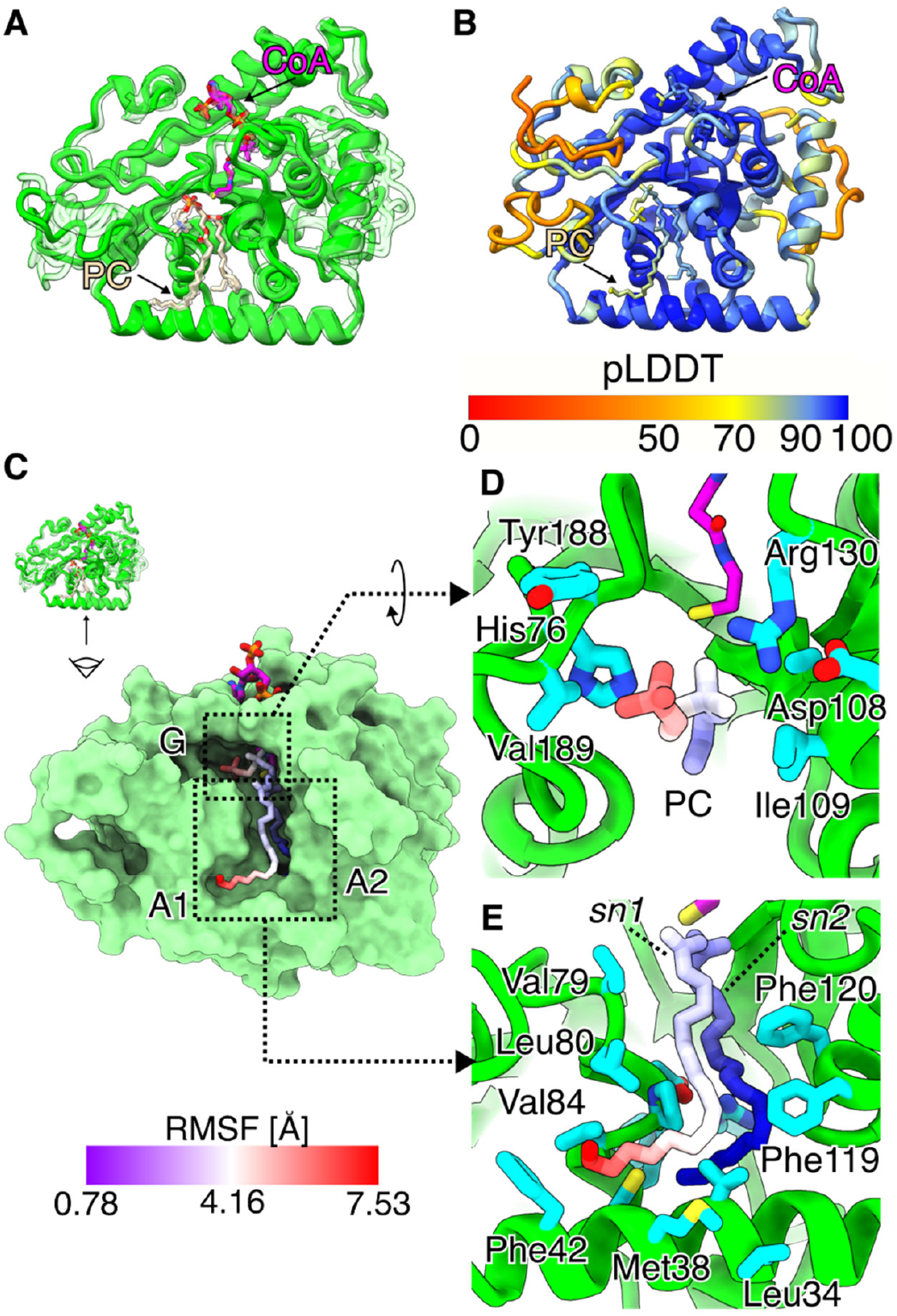
AlphaFold 3 prediction of a complex between tafazzin, PC, and CoA. (A) Superposition of the 5 structures with the highest ipTM scores (transparent) predicted by AlphaFold 3, aligned to the experimental conformation (solid). (B) Local pLDDT scores for the prediction with the highest ipTM. (C) Binding sites and conformational ensemble sampled from simulations. (D) Structure of the G site showing the binding of the glycerol backbone and choline head group of PC. (E) Conformations of the acyl chains of PC bound to the hydrophobic tunnel and the A1 and A2 sites. Atoms of PC are colored by RMSF values.

In summary, it can be concluded that a donor lipid can form a productive enzyme-substrate complex with tafazzin, but only if the central binding site for the acyl chain is unoccupied. This means that simultaneous binding of donor lipid and acyl-CoA into the active site is structurally not feasible.

### Transfer of an acyl chain between two simultaneously bound lipid molecules is highly unlikely

So far, we have only considered the binding of one lipid or lysolipid in the active site of the enzyme together with acyl-CoA or CoA, respectively. In the past, under the assumption that CoA is not involved, it has been speculated that tafazzin can bind a donor lipid and an acceptor lipid simultaneously and that transesterification takes place without intermediate binding of the acyl chain to the protein (*6*). It seems conceivable that in such a mechanism, bound CoA could be responsible for the transfer of the acyl chain in a transition state with two simultaneously bound lipid species. We thus asked the question whether it is possible to bind MLCL and PC to tafazzin in a configuration that would permit direct transfer of the acyl chain.

AlphaFold 3 predictions of the protein with both MLCL and PC reveal that, despite finding binding cavities for both lipids, none of the models place them simultaneously close to the catalytic site (Fig. 6A-C). When we used CL and lysoPC, AlphaFold 3 predicted CL bound to the catalytic site while lysoPC was found in alternative binding sites (Fig. 6D-F), without direct contacts between lipids. Only when acyl-CoA is included in the calculation, the lysolipid is consistently predicted bound to the catalytic site (Fig. 6G,H). The calculated contact probabilities between the hydroxyl group in MLCL and the carbonyl carbon atoms in the acyl chains bound to PC are thus close to zero (Fig. 6I). Interestingly, in the two-lipid predictions, MLCL is bound in conformations similar to those observed experimentally at the tafazzin-ND2/NDUFC2 interface (fig. S8), increasing the confidence in AlphaFold 3 to identify lipid binding sites. We therefore conclude that lipid and lysolipid molecules must bind sequentially to tafazzin, and direct transfer of the acyl chain can be excluded. In such a mechanism, CoA would function as an enzyme bound carrier of the acyl chain while lipid substrates are exchanged.

**Figure 6.**
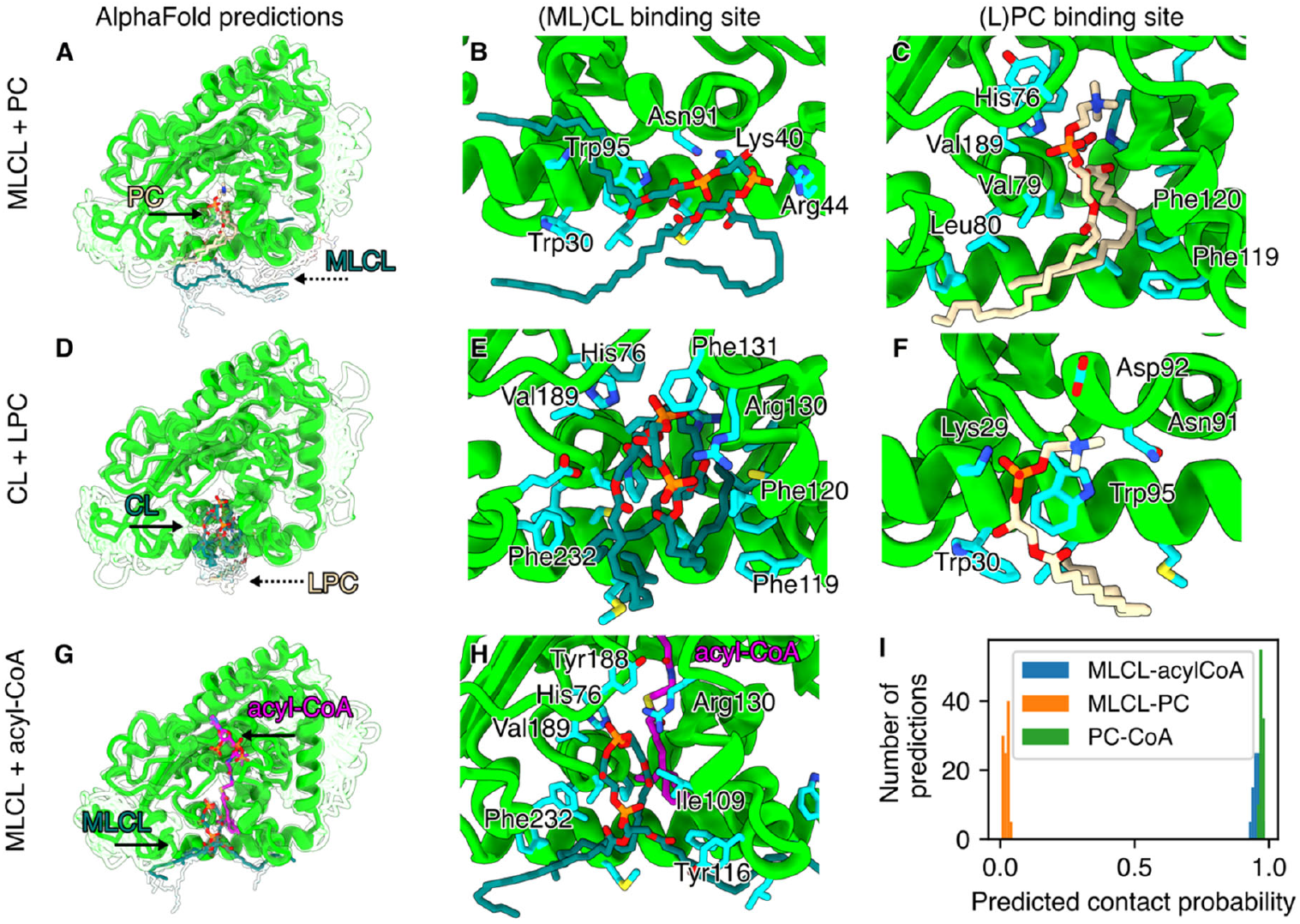
AlphaFold 3 predicts that the formation of a ternary complex between phospholipids and tafazzin is unlikely. (A) Predictions generated for tafazzin in complex with both PC and MLCL. Dashed arrows indicate that the lipid is bound behind the protein with respect to the view. Continuous arrows indicate that the lipid is bound in front of the protein. (B-C) Binding sites of the highest scored predictions of MLCL and PC, respectively. (D) Predictions generated for tafazzin in complex with CL and lysoPC (LPC). (E-F) Binding sites of the highest-scored predictions of CL and LPC, respectively. (G) Predictions generated for tafazzin in complex with MLC and acyl-CoA. (H) Binding site of the highest-scored prediction of MLCL. (I) Predicted contact probabilities between the reactive groups (hydroxyl in MLCL, carbonyl in PC, and thioester in acyl-CoA) for every prediction. AlphaFold 3 defines the contact probability as the probability that the distance between two tokens (residues in proteins and heavy atoms in ligands) will be smaller than 8 Å.

### AlphaFold 3 predicts consistent acyl-CoA and MLCL binding in yeast and human tafazzin

After confirming a strong correlation between the acyl-CoA binding sites in our cryo-EM structure and the AlphaFold 3 model for tafazzin from *Y. lipolytica*, we sought to verify whether human tafazzin also possesses binding sites for acyl-CoA and a lyso-phospholipid at a distance suitable for catalysis. We therefore used AlphaFold 3 to predict a structure for the ternary complex of human tafazzin, MLCL, and linoleoyl-CoA. Tafazzin from humans and yeast have 22 % sequence identity and their AlphaFold 3 models are highly similar, except for short extensions at the N- and C-termini (fig. S9). Strikingly, the predicted binding sites for CoA match closely and show extensive similarities. In the model for the human enzyme, the adenine ring of CoA is also inserted into a distinct binding pocket, one side of which is formed by the indole ring of the conserved Trp164 (*Y. lipolytica* Trp205). The diphosphate group is in contact with Lys163, which corresponds to Arg204 of the yeast enzyme. Similar to the *Y. lipolytica* enzyme, the diphosphate is lined by a glycine-rich sequence stretch that is followed by strictly conserved Gln129 (Gln138 in *Y. lipolytica*). The pantothenic acid moiety is located in a central binding pocket, which enables several interactions with the protein backbone. The methyl groups point towards Val121 and Met132, which correspond to Ile128 and Met141 in the yeast enzyme. It is noteworthy that the sulfur of cysteamine is again positioned in close proximity to the strictly conserved histidine of the active site. Likewise, the interactions of the acyl chain with the three phenylalanine residues described for the yeast enzyme (Phe119, Phe120, Phe184) are also evident in the model for the human enzyme (Phe112, Phe113, Phe148). As in the yeast enzyme, the acyl chain is bent at position Δ-9, with conserved residues Val28, Pro76, and the ion pair Asp75-Arg94 (yeast Val35, Pro83, Asp82, Arg101) located near the bend. Also, the end of the binding pocket is formed by an arrangement of hydrophobic residues that is highly similar to the yeast enzyme. For MLCL, AlphaFold 3 predicts a near perfect match between human and yeast tafazzin with near identical distances between the free hydroxyl group of PA1 and the sulfur atom of acyl-CoA and binding of the PA1 phosphate group by strictly conserved arginine residue (*Y. lipolytica* Arg 130, human Arg123). Furthermore, the narrowest point in the cleft sandwiching the central glycerol and the PA2 glycerol of MLCL is consistently formed by conserved valine (yeast Val189, human Val153) and isoleucine residues (yeast Ile109, human Ile102).

In summary, the conservation of critical residues and AlphaFold 3 structure prediction strongly suggest that acyl-CoA and MLCL bind to human tafazzin in a way that is consistent with the cryo-EM structure and the AlphaFold 3 structure prediction for the yeast enzyme.

### Comparison of acyl-CoA binding sites within the AGPAT family

Given the good match for the acyl-CoA binding sites in tafazzin from yeast and humans, we wondered whether these binding sites might occur in a similar form in other members of the AGPAT family. Unfortunately, only few other structures from this family have been experimentally determined so far (*25, 26, 29, 30*). The cryo-EM structure of human GPAT1 is especially interesting, because it shows the binding pocket of a non-hydrolysable derivative of acyl-CoA (2-oxohexadecyl-CoA) (*26*). Overall, GPAT1 consists of two large domains, with the N-terminal domain exhibiting a typical acyltransferase fold that shows good agreement to the core fold of tafazzin. The positions of the binding pockets for the central parts of acyl-CoA are comparable for both proteins (fig. S10A) and in both cases, the sulfur atoms of the CoA molecules are located near the active site histidines. However, there is only limited conservation of residues lining the pocket and the 3’-phospho-adenosine head groups are located in different positions in binding pockets with distinctly different structures. Instead of the prominent stacking interaction with a tryptophane residue in tafazzin, a stronger involvement of positively charged residues is noticeable in GPAT1. The bacterial lysolipid acyltransferase PlsC is another member of the AGPAT family for which detailed structural information is available (*25*). The overall folds of tafazzin and PlsC are similar (*21*) and MD simulations showed that the sulfur of the acyl donor is situated in close distance to the active site histidine residue (*25*). However, since the acyl-donor for PlsC is an ACP instead of CoA, we employed an AlphaFold 3 model of human LPLAT1 with its substrates for further investigation. LPLAT1 is a canonical member of the AGPAT family that is responsible for the synthesis of phosphatidic acid and is expressed in all human tissues (*8*). AlphaFold 3 models of tafazzin and LPLAT1 show good agreement, but the latter has a putative transmembrane helix and two amphipathic surface helices instead of only one in tafazzin (fig. S10B). Similar to GPAT1, the 3’-phosphoadenosine head group sits in a different position in LPLAT1 than in tafazzin. Trp205, which in tafazzin sits parallel to the adenosine, is replaced by Arg or Lys in LPLAT1 (fig. S4). The CoA binding sites start to converge at the position of the diphosphates, and the pantothenic acid and cysteamine groups, as well as a significant portion of the acyl chain, are bound in the same position in both proteins (fig. S10). However, the prominent bend of the acyl chain at position Δ-9 observed in tafazzin is absent in LPLAT1, consistent with the replacement of Pro83 and Asp82 by hydrophobic residues, the absence of the loop containing Arg101, and replacement of Phe119 and Phe120 by smaller hydrophobic residues (fig. S4). It is also interesting to note that the phosphate of lyso-PA in LPLAT1 sits in the same position as the PA1 phosphate of MLCL and is bound by a corresponding arginine residue. In summary, it can be concluded that the positions of binding pockets for pantothenic acid, cysteamine, and at least the first C atoms of the acyl chain overlap in the representatives of the AGPAT family examined. By contrast, there are clear differences in both the position and structure of the respective binding pockets for the 3’-phosphoadenosine head groups.

## Discussion

Lipid remodeling is an important cellular process that was first described by Lands as a deacylation-reacylation cycle (*31*). The two complementary reactions are carried out by phospholipases and LPLATs, respectively. Variations of the Lands cycle based exclusively on the forward and reverse reactions of LPLATs have been proposed (*32, 33*). However, all transacylation reactions in the Lands cycle and its variants have in common that the donor for the acylation of transiently formed lysolipids is acyl-CoA. In stark contrast, it has been firmly established that the reaction of the mitochondrial transacylase tafazzin is independent of CoA or CoA derivatives as donors or acceptors of the transferred acyl chain (*7*). Therefore, our discovery that tafazzin from the yeast *Y. lipolytica* binds an acyl-CoA molecule was entirely unexpected. Nevertheless, the evidence from our cryo-EM data is compelling and is supported by high-confidence AlphaFold 3 structure predictions and MD simulations. Furthermore, important residues involved in acyl-CoA binding are conserved, and computational analyses yielded consistent results for human and yeast tafazzin. We therefore conclude that binding of acyl-CoA is a general feature of tafazzin. This is consistent with tafazzin belonging to an enzyme family whose other members all interact with acyl-CoA or acyl-ACP (*8, 9*). In fact, we show that binding of acyl-CoA has some structural similarities with acyl-CoA binding to GPAT1 and LPLAT1. To explore the yet unresolved catalytic mechanism of tafazzin, we analyzed the arrangement of substrates in the active site by complementing our structural data with structure predictions and MD simulations.

We consistently found that the sulfur atom forming the thioester bond in acyl-CoA is close to the active site histidine and the appended fatty acid is inserted into a binding cavity suitable to accommodate a C18 acyl chain. The kink of the acyl chain at the position of the *cis* double bond at position Δ-9 and the interaction of phenylalanine residues with double bonds seen in MD simulations suggest a preference of tafazzin for at least Δ-9 unsaturated fatty acids. The guanidinium group of Arg101 of the conserved Asp-Arg ion pair might be involved in a cation-π pair with additional double bonds of the acyl chain. The finding of higher transacylation activities for unsaturated acyl chains in biochemical assays (*7, 10, 11*) is thus in full agreement with our data. However, several lines of evidence indicated that the inherent preference of tafazzin for acyl chains plays only a limited role. Transgenic expression of human tafazzin in *Saccharomyces cerevisiae* (*34*) or in *Drosophila melanogaster* (*35*) reproduced the native CL acyl chain composition of the yeast or the fly, respectively. Furthermore, it was shown that the observed variations in the lipid tail composition of CL in different human tissues did not correlate with the expression profiles of the tafazzin gene (*36*). Instead, it has been proposed that the availability of specific fatty acids in the phospholipid pool is important (*36*) and that the physical properties of the membrane determine which types of lipids are formed in tafazzin-mediated transacylation reactions (*12*). This is consistent with the fact that acyl-CoA can bind even with unsaturated acyl chains, though without benefit of the cation-π interactions described above.

The formation of functionally important CL species, including the well-known (18:2)_4_ CL of the heart, requires that the free hydroxyl group of PA1 in MLCL be reactive in both the *sn-*1 and *sn-*2 positions. Our MD simulation data show that both variants of MLCL bind appropriately to the enzyme to allow interaction of their free hydroxyl groups with the histidine residue of the active site. The abstraction of a proton enables a nucleophilic attack on the thioester bond of acyl-CoA, which is located at a suitable distance. The resulting transition state is likely stabilized by an oxyanion hole involving the backbone of Gly105 and Ala106. We therefore conclude that the acyl chain of a bound acyl-CoA can be transferred to a lyso-phospholipid following a similar mechanism as previously proposed for PlsC (*25*) and GPAT1 (*26*) (Fig. 7).

**Figure 7.**
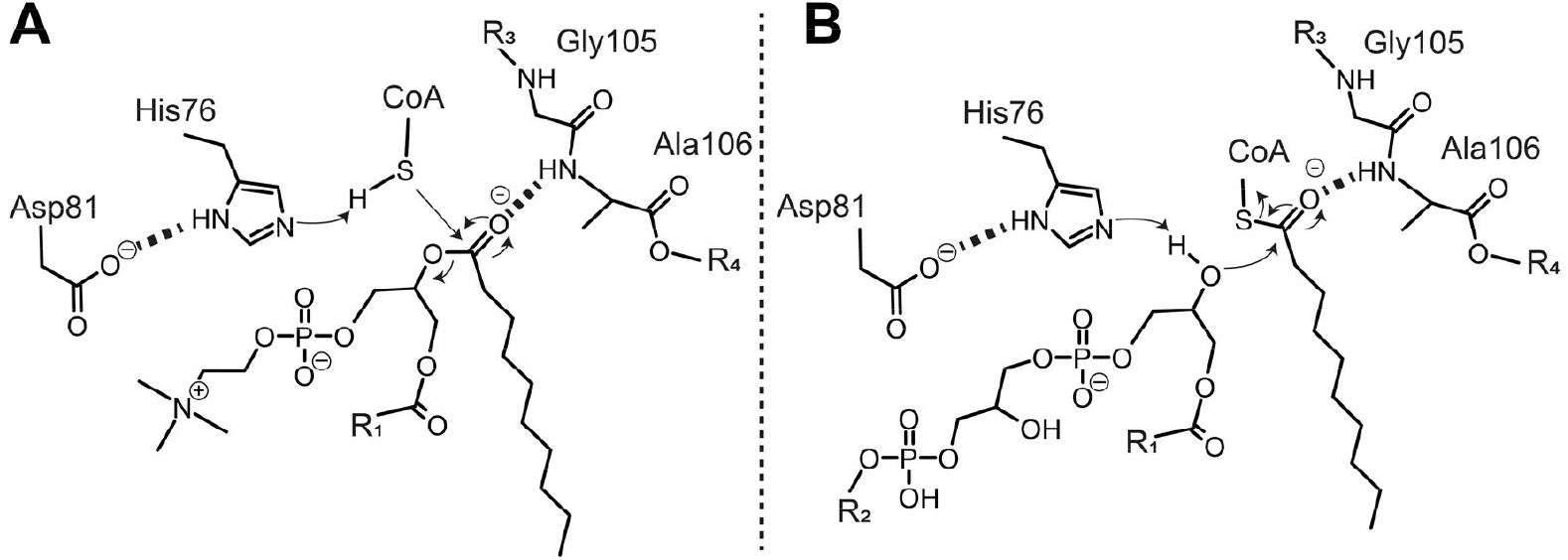
Mechanism of transacylation. (A) Transfer of an acyl chain from PC to CoA. His76 abstracts a proton from the thiol of CoA to enable a nucleophilic attack on the *sn2* ester bond of PC. The oxyanion intermediate is stabilized by the Gly105/Ala106 oxyanion hole. (B) Transfer of an acyl chain from acyl-CoA to MLCL. His76 abstracts a proton from the free hydroxyl of MLCL (shown here for *sn2*), enabling a nucleophilic attack on the thioester bond of acyl-CoA. The oxyanion intermediate is stabilized by the Gly105/Ala106 peptide bond. R_1_: acyl chain; R_2_: diacylglycerol; R_3_: tafazzin amino acid 104; R_4_: tafazzin amino acid 107.

Given that tafazzin is a transacylase, the origin of the transferred acyl chain must be a lipid molecule. Structure prediction and MD simulations indicated that in a complex of PC, CoA, and tafazzin, the *sn*-2 acyl chain inserts into the lipid binding cavity and the ester bond to be cleaved is in close vicinity of the thiol group of CoA. In fact, a preference of *sn*-2 over *sn-*1 for the position of the acyl chain to be abstracted from the donor lipid has been reported previously based on biochemical data (*10, 11*). Again, the backbone of Gly105 and Ala106 is in suitable distance to stabilize the transition state. We therefore propose that CoA bound to tafazzin receives and stores an acyl chain from a donor lipid, employing a mechanism where the thiol group of CoA is activated by abstraction of its proton by the active site histidine followed by a nucleophilic attack on the *sn-*2 ester bond (Fig. 7). The conversion of an oxo-ester into a thio-ester does not pose a principal problem and was shown, e.g., in the activation steps of fatty acid synthase where an acyl group is transferred from a serine residue to the thiol group of the ACP (*37*) and in the reverse reaction of LPLATs (*32, 33*).

Regarding the mechanism of tafazzin, we also consider it important that consecutive binding of donor lipids in the active site is not possible, since the acyl chain binding cavity is occupied by an acyl chain that was transferred to CoA in the previous reaction cycle. If acyl-CoA remains bound to the enzyme, the site only becomes accessible again after the acyl chain has been transferred to a lysolipid, preventing uncontrolled repetition of deacylation reactions. It follows that the presence of lysolipids is a prerequisite for the activity of tafazzin, which is consistent with the biochemical data. It has been established that MLCL is formed by the phospholipase Cld1 in yeast (*38*) and it was recently shown that ABHD18 is responsible for MLCL formation in humans (*39*). Schlame and colleagues proposed that tafazzin simultaneously binds a phospholipid and a lyso-phospholipid in a ternary complex, forming an enzyme-substrate complex where two lysolipids share an uncommitted acyl chain between them (*6*). One could envisage that in such a mechanism CoA could serve as a transient carrier of the acyl chain. However, our MD simulations argue strongly against simultaneous binding of PC and MLCL with suitable distances of functional groups in the active site. We thus propose that the binding of lipid substrates occurs sequentially and that CoA is an intermediate enzyme-bound carrier of the acyl chain while lipid substrate molecules are exchanged. Such a function implies that either loaded or unloaded CoA is present in the CoA binding site of tafazzin and that CoA has more of the characteristics of a prosthetic group than those of a coenzyme. Covalent binding of a modified CoA as a prosthetic group has been reported for selected bacterial enzymes (*40, 41*), but is unknown for eukaryotic enzymes. We don’t observe a covalent interaction of CoA with tafazzin; however, it is striking that the binding of the 3’-phospho-adenosine group of CoA occurs in a pocket that is different from that of GPAT1 and LPLAT1 and where a strong stacking interaction with a highly conserved tryptophan residue can form. MD simulations show a persistent binding of CoA in its binding pocket and the strong density in our cryo-EM map for CoA also indicates a high occupancy of the CoA binding site. The weaker density for the acyl chain suggests that in our preparation, a fraction of the tafazzin molecules have unloaded CoA bound, in agreement with the model proposed above. Since we did not add CoA or CoA derivatives at any step of protein purification or sample preparation, we are confident that our structure faithfully reflects the status of tafazzin in native membranes.

Our new structural data and insights into the mechanism of transacylation now enable a molecular understanding of Barth syndrome mutations. We classify missense mutants listed in the Human *Tafazzin* Gene Variants Database (*42*) into five groups (Fig. 8, Table S1, movie 5). Several exchanges of corresponding amino acid residues have already been introduced into tafazzin from *S. cerevisiae* by site-directed mutagenesis, and their effects on expression and activity have been analyzed, but in absence of structural information a molecular understanding was not possible at the time (*20*). In this discussion, we will use exclusively the numbering for the human enzyme (excluding exon 5 (*34*)), but Table S1 allows for a translation of the residue numbers between tafazzin from humans, *Y. lipolytica*, and *S. cerevisiae*. Mutants in **group 1** affect residues in or near the active site. These include the His69Gln and Asp74Glu mutations of the HisX_4_Asp motif. The mutations Ser71Pro, His146Tyr/Arg/Pro, Phe148Ile/Leu, Val153Gly and His184Arg affect residues located nearby. Residues Phe148 and Val153 sandwich the active site histidine. Interestingly, the phenylalanine is also involved in the binding of the acyl chain attached to CoA and the valine contributes to the binding pocket of MLCL. Corresponding mutations in tafazzin from *S. cerevisiae* caused loss of transacylase activity (*20*). The residue His146 is highly conserved and points to the aromatic ring of Phe148. However, the functional role and the reason for its conservation in tafazzin are still unclear. His184 is located at hydrogen bonding distance from Asp74 of the active site. The mutations in **group 2** affect residues involved in the binding pocket for the acyl chain to be transferred to or from CoA. These include Val28, Pro76, and Gly116, all of which are located near the hinge point at position Δ-9 of the acyl chain. A mutation corresponding to Gly116Asp caused loss of activity of the *S. cerevisiae* enzyme. Cys118 is located in the section of the binding pocket upstream of the hinge point and an exchange with a larger residue likely interferes with binding of the acyl chain. As mentioned above, Phe148 can be assigned to both group 1 and group 2, as it is intimately associated with the active site His69 and is located near the first methylene units of the acyl chain close to the thioester bond. An exchange of a tyrosine residue at the corresponding position results in activity loss of tafazzin from *S. cerevisiae*. It is interesting to note that the ion pair formed by Asp75 and Arg94 is a hot spot for pathogenic mutations. We speculate that the guanidinium group of the Arg residue interacts with double bonds in the acyl chain. In any case, mutations in these two residues are likely to disturb the structure of the acyl chain binding pocket. **Group 3** combines mutations that affect residues in or near the CoA binding pocket. These include Gly124Arg and Gln129Pro, which are located in the binding motif for the diphosphate. Gln129 is highly conserved and makes contacts with the backbone oxygen and nitrogen of position 122, thereby stabilizing the loop interacting with the diphosphate. A larger residue at the position of Gly165, i.e., as in mutant Gly165Val, interferes with the binding of pantothenic acid. A hot spot for Barth syndrome mutations, Gly167, is located only two positions away. Finally, Thr273 sits in the binding pocket for the adenine ring and the Thr237Lys mutation interferes sterically with binding of CoA. **Group 4** contains residues Val153 and Cys103 which are both involved in binding of substrate lipid molecules. The pathogenic exchange Val153Gly was tested in *S. cerevisiae* tafazzin and had a clear impact on transacylase activity. The link of mutant Cys103Arg with Barth syndrome is less clear because it was found in a patient carrying an additional mutation in myosin-7. However, its position in the structure strongly suggests that the function of tafazzin is compromised because the larger side chain of arginine would block the acyl chain binding pocket. Mutants of Asp101 are also added to group 4; however, lipid binding is only indirect by interacting with Arg123 that is critical for binding the phosphate of the lipid head group. In **group 5** we list mutations with less clear impact on tafazzin function. For most of them, destabilization of the structure is plausible. It is remarkable that six affected hydrophobic residues are clustered in a single beta strand (position 178 to 185) and in an adjacent helix (position 47 to 56).

**Figure 8.**
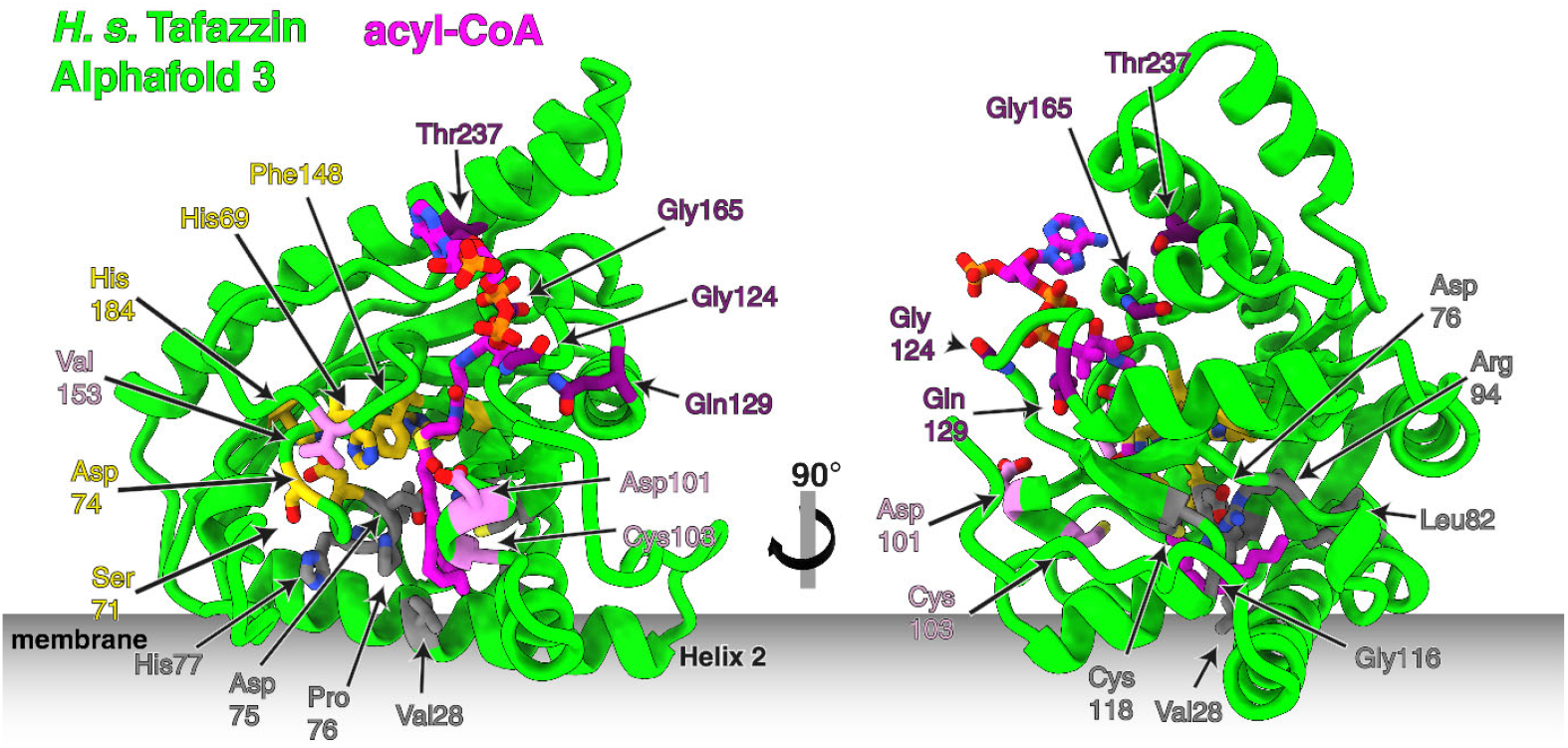
Positions in human tafazzin with known Barth syndrome mutations. Residues are colored according to classification in group 1 (yellow), group 2 (light grey), group 3 (purple), group 4 (violet); details see text and Table S1. Group 5 comprises mutations with less clear impact on tafazzin function. The model was generated by AlphaFold 3 with acyl-CoA as a ligand from the *H. sapiens* sequence missing exon 5.

Taken together, our work provides detailed insights into the transacylation mechanism of tafazzin and offers a molecular explanation of how pathogenic mutations cause tafazzin dysfunction. The most surprising finding is that tafazzin binds acyl-CoA in a catalytically competent manner. Given that biochemical data have so far excluded CoA or acyl-CoA as an exchangeable substrate, we suggest that CoA in tafazzin has the function of a prosthetic group to transiently bind an acyl chain exchanged between a lipid/lysolipid pair.

## Methods

### Purification of complex I assembly intermediates

Cell growth, preparation of mitochondria and protein purification was performed as described previously (*21*), but using lauryl maltoside neopentyl glycol (LMNG) at a ratio of detergent to mitochondrial protein of 0.8 g/g for solubilization and at a concentration of 0.002 % in all buffers during subsequent chromatography.

### Sample vitrification and cryo-EM data acquisition

The samples were vitrified by rapid freezing in liquid ethane using an FEI Vitrobot Mark IV. Three microliters of protein at a concentration of 1.3 mg/ml were applied to Quantifoil 1.2/1.3 300-mesh grids (Quantifoil Micro Tools GmbH), which had been glow-discharged in a PELCO easiGlow system at 15 mA for 90 s. The freezing conditions were set to: chamber temperature of 4 °C, 100% relative humidity, blot force of 3 (Whatman 595), and blot time of 5 s. Data was automatically recorded via the EPU software (Thermo Fisher Scientific) on a Krios G4 cryo-TEM (Thermo Fisher Scientific) at 300 kV. The Krios G4 was equipped with an E-CFEG, a Selectris X energy filter, and a Falcon 4i direct electron detector. A total of 13,250 movies were recorded with a defocus ranging from -0.8 to -1.8 µM. The electron filter was set to a slit width of 10 eV, and a 50-µM aperture was used for data acquisition. The movies were recorded in EER format at a dose of 55 e/Å^2^. The nominal magnification was set to 215,000x, resulting in a calibrated pixel size of 0.573 Å.

### Cryo-EM data processing

RELION5 was used for data processing (*43, 44*). CryoSPARC4.4.0 was used for initial data assessment and ab initio model generation. Final maps were subjected to density modification using Phenix refine (*45*).13,252 movies were imported to RELION subdivided into 32 EER fractions . Motion-correction was performed applying the MotionCor2 algorithm (*46*). Motion-corrected micrographs were processed using the CTFfind4 implementation of RELION, to estimate contrast transfer function (CTF) parameters (*47*). Particles were picked with Topaz (*48*). Initially, 1,874,208 particles were picked and assessed by 2D classification. The worst classes were discarded, and the rest of the particles were imported into cryoSPARC and used for an ab initio reconstruction (*44*). The ab initio reconstruction was done with three final structures, allowing for sorting of different classes in 3D while simultaneously generating a density map. Two classes only yielded a noise map, while one showed a distinct particle shape. This class was refined in a non-uniform refinement job and the resulting map was used as a template for 3D classification of the picked particles in RELION. After several rounds of 3D classification, 389,400 good particles were used for 3D refinement. An iterative approach of CTFrefine and map refinement was used until a final resolution of 2.3 Å was reached. This map was further sharpened by phenix density modification to a final resolution of 2.2 Å (*45*). Local resolution was estimated by phenix.local_resolution (*49*).

### Model building

Model building was done in Coot (*50*) using the published structure PDB: 7ZKQ as a starting model. The structure was refined using phenix real space refinement and assessed with molprobity (*51, 52*).

### AlphaFold 3 predictions

The sequences of *Yarrowia lipolytica* and *Homo sapiens* tafazzin were retrieved from UniProt with the accession codes Q6CBZ7 and Q16635, respectively. Molecular structures of acyl-CoA, CoA, phosphatidyl choline, and monolysocardiolipin with a reference conformation were prepared in cif format to be used for predictions. A local installation of AlphaFold 3 was run to predict the structure of the proteins in complex with the ligands. We requested 5 samples from 20 different random seeds, yielding a total of 100 predictions per complex. The predicted complexes were tafazzin (either *Y. lipolytica* or *H. sapiens*) with linoleoyl-CoA and tri-palmitoyl-*sn*2-hydroxyl MLCL, and CoA with *sn*1-stearyl-*sn*2-linoleoyl phosphatidyl choline. The resulting predictions were sorted by ipTM score and the top-ranked structures were selected as starting conformations for MD simulations. The local pLDDT scores were also visualized for the top-ranked structures using ChimeraX.

Additionally, the sequence of *H. sapiens* lysophospholipid acyltransferase 1 (LPLAT1, following Valentine’s nomenclature (*8, 9*)) was retrieved from the UniProt accession code Q99943. AlphaFold 3 predictions of this protein in complex with acyl-CoA and lysophosphatidic acid were prepared following the same protocol described above.

### Molecular dynamics simulations

Selected AlphaFold 3 predictions were used in the Membrane Builder tool of CHARMM-GUI (*53, 54*) to generate a simulation box containing tafazzin complexes at an explicit membrane in aqueous solution at physiological salt concentration. The composition of the membrane was 50% *sn*1-stearyl-*sn*2-linoleoyl phosphatidyl choline (SLPC), 30% *sn*1-palmitoyl-*sn*2-oleoyl phosphatidyl ethanolamine (POPE), and 20% tetralinoleoyl cardiolipin (TLCL1), as commonly used to model the inner mitochondrial membrane (*55, 56*). The CHARMM36m force field (*57*) was used to model the protein and the CHARMM36 lipid parameters were used for the membrane. Water was modeled using the CHARMM-compatible TIP3P parameters. Force field parameters for MLCL were adapted from CL topologies as described in (*58*). Parametrization of (acyl-)CoA was done following a fragmentation approach. Parameters from the RNA CHARMM force field were used for the adenosine group of (acyl-)CoA. The pantothenic-cysteamine scaffold was modelled using bonded and Lennard-Jones parameters from the CHARMM general force field (CGenFF) (*59*). Partial charges on the pantothenic-cysteamine fragment were calculated using electronic structure methods as follows: conformers of the fragment were generated using the global geometry optimization and ensemble generator (GOAT) (*60*) algorithm in Orca 6.0.1 (*61*) with the XTB2 semiempirical (*62*) to accelerate calculations. Low energy conformers were used in single-point energy calculations with the MP2/6-31G(d) level of theory, as recommended for CHARMM (*59*), and implicit polarization with the CPCM model, using water as solvent. For every conformer, the electrostatic potential around a Merz-Kollman grid was calculated. Classical partial charges were calculated by fitting the quantum electrostatic potential using the standard 2-stage RESP method (*63*).

Molecular dynamics simulations were performed in Gromacs 2025.1 (*64*). The initial systems were first energy-minimized with the steepest descent method. After energy minimization, a modified equilibration protocol based on the standard protocol of CHARMM-GUI for membrane systems was followed as listed in Table 1.

**Table 1.** Equilibration protocol followed in simulations. Six simulations (top to bottom) were performed with decreasing restraints. The force constant of the backbone atoms was also applied to heavy atoms of (acyl)-CoA and the bound phospholipid. Dihedral restraints were applied to acyl chains of every phospholipid double bond to enforce *cis* stereochemistry. The last three equilibration steps were simulated with distance restraints between the protein and both ligands (acylCoA and bound phospholipids) with the force constant reported in the Backbone column.

| Integration time step [fs] | Ensemble | Simulation length [ns] | Position restraints force constant [kJ mol<sup>-1</sup> nm<sup>-2</sup>] |  |  |  |
| --- | --- | --- | --- | --- | --- | --- |
|  |  |  | Backbone | Side chains | Phospholipid phosphate | Acyl chain dihedral |
| 1 | NVT | 1 | 4000 | 2000 | 1000 | 1000 |
| 1 | NVT | 1 | 2000 | 1000 | 400 | 400 |
| 1 | NPT | 1 | 1000 | 500 | 400 | 200 |
| 2 | NPT | 5 | 500 | 200 | 200 | 200 |
| 2 | NPT | 5 | 200 | 50 | 40 | 100 |
| 2 | NPT | 5 | 50 | 0 | 0 | 0 |
| 2 | NPT | 50 | 500 | 0 | 0 | 0 |
| 2 | NPT | 50 | 250 | 0 | 0 | 0 |
| 2 | NPT | 50 | 100 | 0 | 0 | 0 |

After equilibration, each system was simulated for 1 μs, with three independent replicates per system. Simulations were done in the NPT ensemble with a time step of 2fs, using the velocity-rescaling thermostat (*65*) with a target temperature of 310 K and a coupling time of 1.0 ps and the Parrinello-Rahman (*66*) barostat with a target pressure of 1 bar and a coupling time of 5.0 ps. Electrostatic interactions were treated with the particle-mesh Ewald method (*67*) using a real-space cutoff of 12 Å for both Lennard-Jones and electrostatic interactions. All covalent bonds involving hydrogen atoms were kept rigid using the LINCS algorithm (*68*). Finally, the SETTLE algorithm was used to keep water molecules rigid (*69*).

### Trajectory analysis

Trajectories from MD simulations were analyzed using custom Python code based mainly in the MDAnalysis library (*70*) as described below. First, the DSSP tool (*71*) in MDAnalysis was used to calculate an average secondary structure of the protein along trajectories. Then, the folded parts of tafazzin were used to align the trajectory to the first frame of the simulation. Root mean squared fluctuation quantities were calculated using this alignment as reference.

To characterize lipid (or CoA)-protein interactions and identify binding sites, a contact score between every pair of heavy atoms (*c*_*ij*_) was calculated for every frame on the trajectory following equation 1 (*72*),

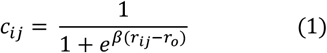

where *r*_*ij*_ is the distance between heavy atoms of the protein and the lipid, *r*_*o*_ is a soft cutoff, and β controls the softness of the switching function. The values of r_o_ = 4.5 A and β=5.0 were selected, as recommended in the MDAnalysis documentation. The per-atom contact score was averaged over trajectories and aggregated by summation over residue atoms to generate a residue-level contact score. Residues forming persistent interactions (average *c*_*i*_ > 1.0) with lipids were identified from the contact score and binding sites were proposed based on this analysis.

## Acknowledgments

We thank the Central Electron Microscopy Facility of the Max Planck Institute of Biophysics headed by Sonja Welsch for cryo-EM infrastructure and technical support; Juan Castillo-Hernández, Özkan Yildiz, the Central IT team and the Max Planck Computing and Data Facility for support in cryo-EM data processing. We thank Karin Siegmund for excellent technical assistance. Fruitful discussions with Michael Schlame (New York University School of Medicine) are gratefully acknowledged. We thank Balázs Fábián and Tobias Hüfner for discussion and feedback on simulations.

## Author contributions

JGRJ prepared AlphaFold calculations, run and analyzed the MD simulations, drew figures, contributed to writing the manuscript; JS carried out the biochemical work, prepared cryo-EM grids, acquired and processed cryo-EM data, built and analyzed atomic models, drew figures, contributed to writing the manuscript; JV built and analyzed atomic models, interpreted the structures, and contributed to writing the manuscript, GH analyzed MD simulations and contributed to writing the manuscript, VZ initiated the project, interpreted the mechanistic implications of the structures, wrote the first draft of the manuscript and drew figures.

## Data and materials availability

Cryo-EM maps and model coordinates will be deposited in the wwPDB. The structures of the AlphaFold predictions calculated in this work are available in Zenodo (10.5281/zenodo.18307766).

