## supplemental information for "Coenzyme A is bound to tafazzin – a paradigm change for transacylation"

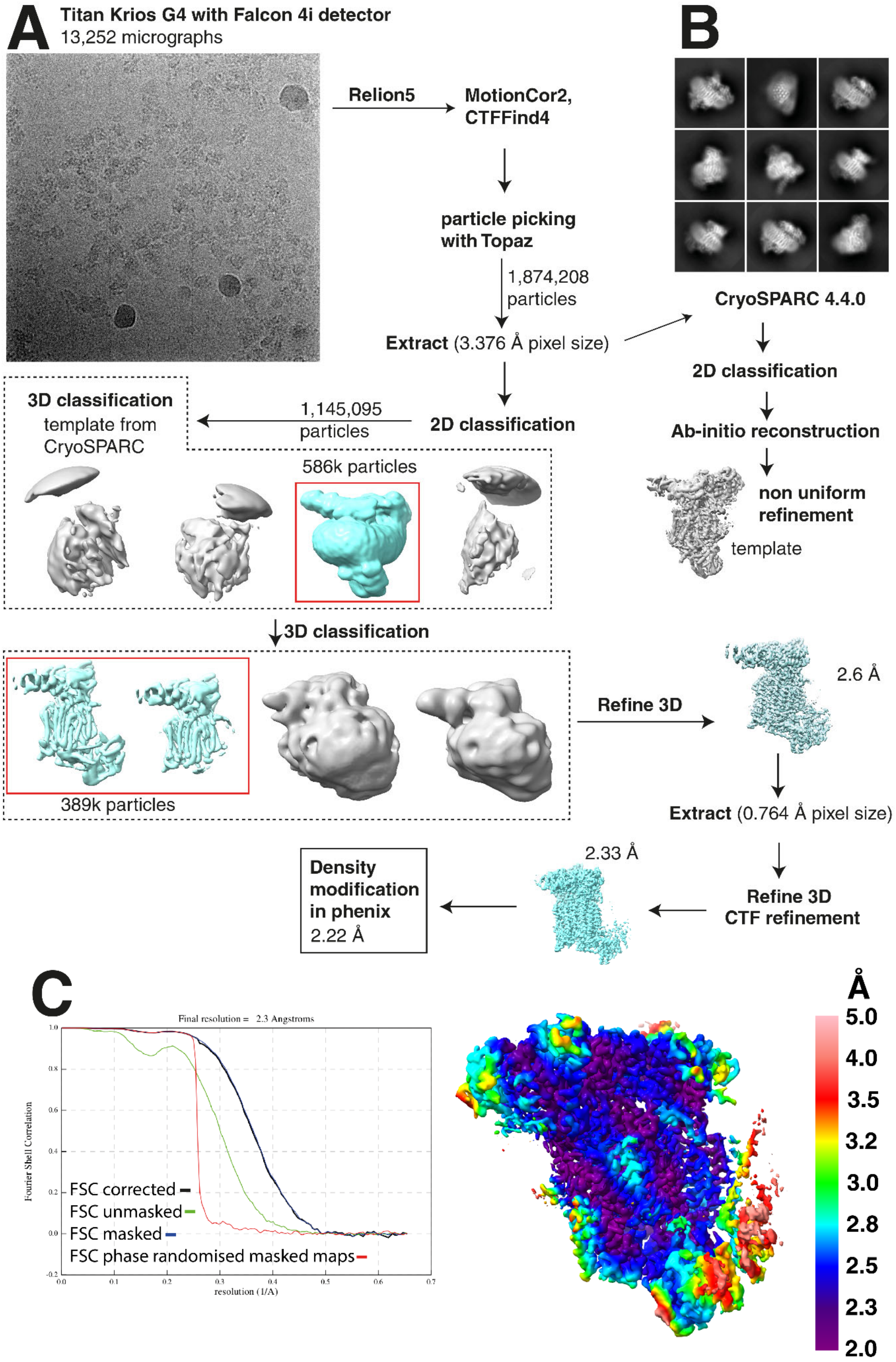

A
Titan Krios G4 with Falcon 4i detector
13,252 micrographs
Relion5
MotionCor2,
CTFFind4
particle picking
with Topaz
1,874,208
particles
Extract (3.376 Å pixel size)
2D classification
1,145,095
particles
3D classification
template from
CryoSPARC
586k particles
3D classification
389k particles
Refine 3D
2.6 Å
Extract (0.764 Å pixel size)
Refine 3D
CTF refinement
2.33 Å
Density
modification
in phenix
2.22 Å
B
CryoSPARC 4.4.0
2D classification
Ab-initio reconstruction
non uniform
refinement
template
C
Final resolution = 2.3 Angstroms
Fourier Shell Correlation
resolution (1/A)
FSC corrected
FSC unmasked
FSC masked
FSC phase randomised masked maps
Å
5.0
4.0
3.5
3.2
3.0
2.8
2.5
2.3
2.0

**Figure S1: Cryo-EM data processing**

(A) Processing of the dataset and a representative micrograph (see methods). (B) representative classes from a 2D classification performed in RELION. Processing was carried out in RELION5, unless otherwise specified. Density modification was performed using phenix.resolve_cryo_em. (C) Gold-standard Fourier shell correlation curve and map colored by local resolution estimated by phenix.local_resolution.

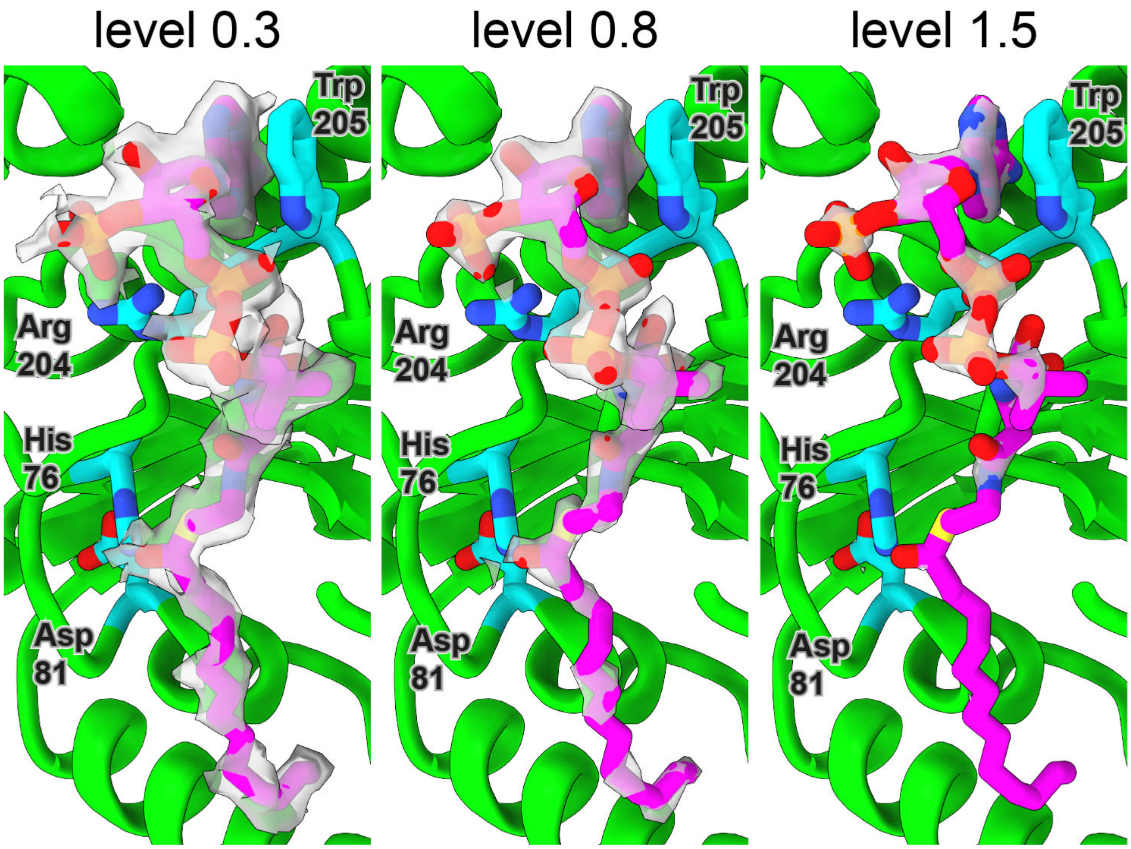


**Figure S2: Cryo-EM density of acyl-CoA bound to tafazzin.** The density within 2 Å distance of the modeled acyl-CoA is depicted at different isocontour levels. The $HisX_4Asp$ motif is in close proximity to the thioester of CoA. Residues Arg204 and Trp205 interact with the phosphates and the adenine of the headgroup, respectively.

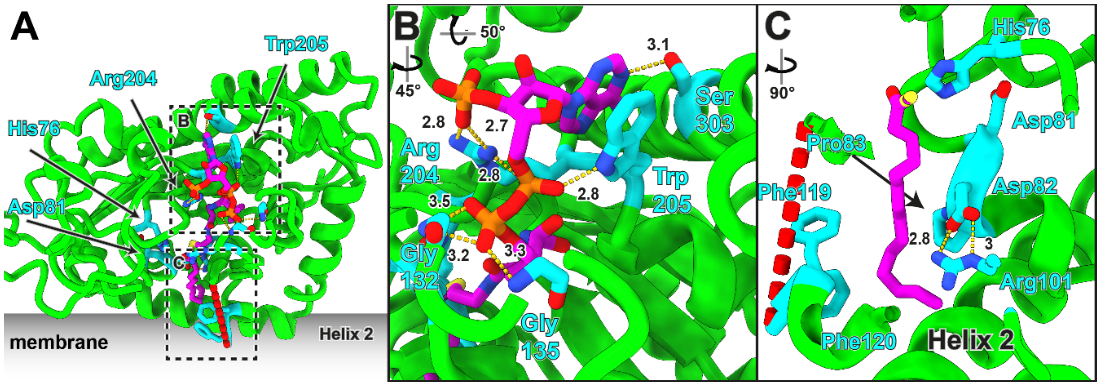


**Figure S3: Coordination of acyl-CoA in tafazzin.** (A) cryo-EM structure of tafazzin with bound acyl-CoA. (B) Detail view of the 3' phosphoadenosine and the diphosphate coordination in tafazzin with distances in Å. (C) The thioester bond of the acyl chain is close to His76. The acyl chain has a curved conformation in its binding pocket. The conserved Asp82/Arg101 salt bridge, that is unique to tafazzin in the AGPAT family, lines the acyl chain cavity. Pro83 forms a *cis* peptide bond and is important to position Asp82 relative to Arg101.

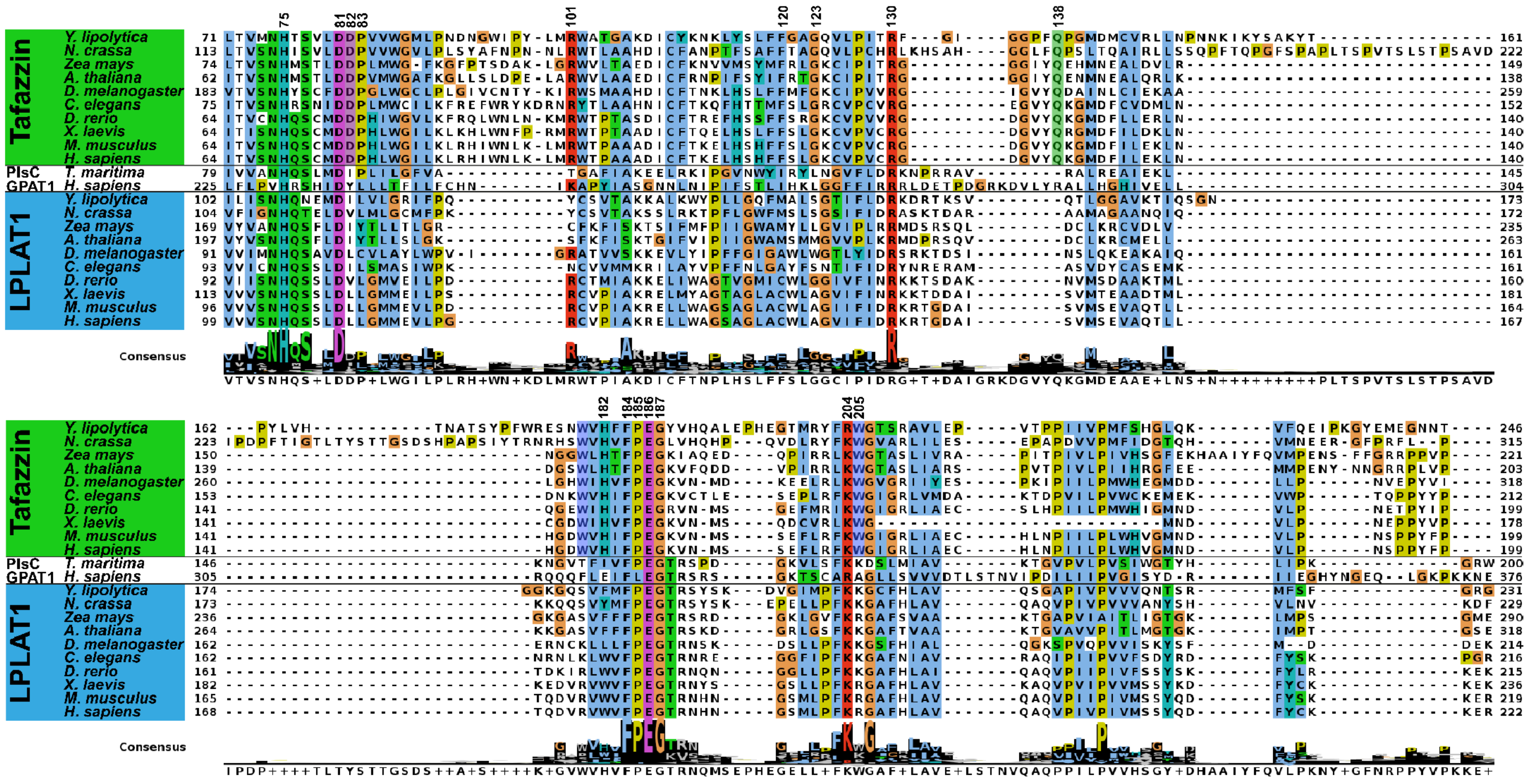
Tafazzin
LPLAT1
PlsC
GPAT1
Y. lipolytica
N. crassa
Zea mays
A. thaliana
D. melanogaster
C. elegans
D. rerio
X. laevis
M. musculus
H. sapiens
T. maritima
Consensus
75
81
82
83
101
120
123
130
138
182
184
185
186
187
204
205

**Figure S4: Sequence alignment of conserved regions between tafazzin and LPLAT1.**

Tafazzin sequences used are: Q6CBZ7 *Y. lipolytica*; Q7S116 *N. crassa*; A0A804RN79 *Z. mays*; Q9ZV87 *A. thaliana*; Q9V6G5 *D. melanogaster*; Q23598 *C. elegans*; F1QCP6 *D. rerio*; Q7ZXM1 *X. laevis*; Q91WF0 *M. musculus*; Q16635 *H. sapiens*. PlsC and GPAT1 sequences Q9X219 *T. maritima* and Q9HCL2 *H. sapiens* were used. LPLAT1 sequences used in the alignment: Q6C5D5 *Y. lipolytica*; Q7RXP0 *N. crassa*, C0PEF9 *Z. mays*, Q8GXU8 *A. thaliana*; Q7KTI0 *D. melanogaster*; Q22267 *C. elegans*; A0JMM2 *D. rerio*; A0A1L8F3L6 *X. laevis*; O35083 *M. musculus*; Q99943 *H. sapiens*. The alignment was generated using the ClustalO algorithm and is visualised using Jalview. Numbers above (on the right) the alignment indicate the residue numbers in *Y. lipolytica*. Table S1 allows for a translation of disease mutant positions between yeast and human.

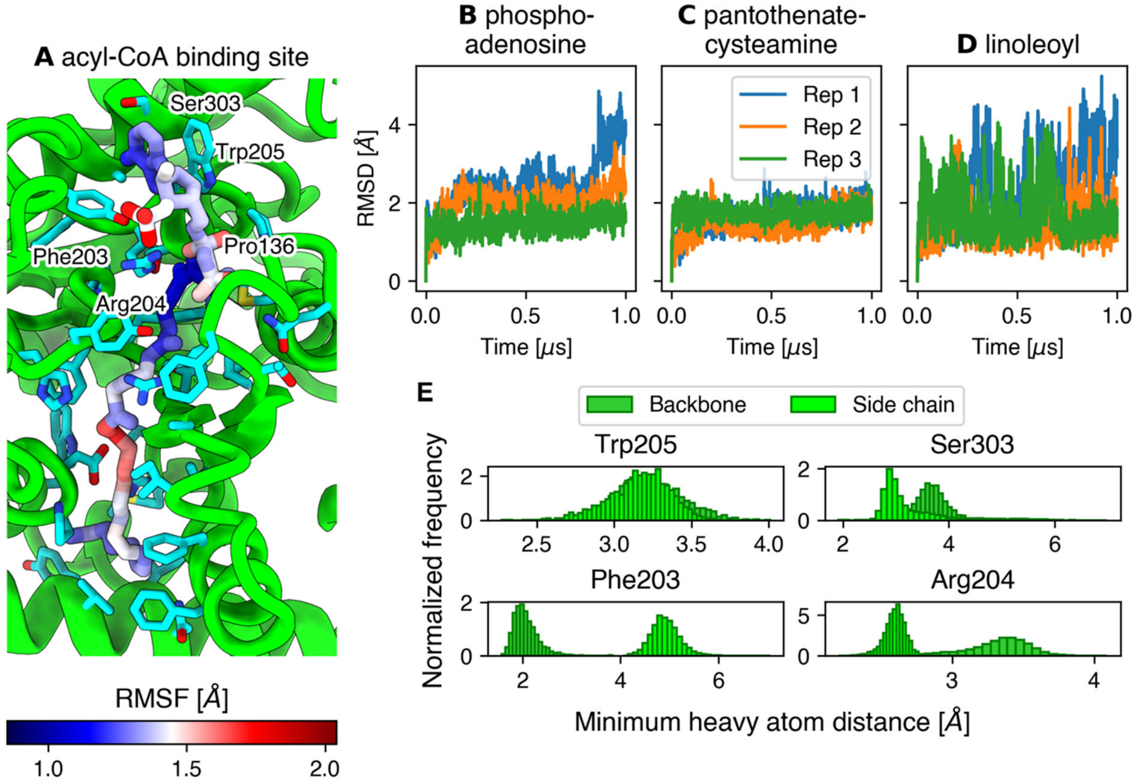


**Figure S5. Acyl-CoA binding mode in MD simulations.** (A) Conformation of acyl-CoA bound to tafazzin observed in simulations. Acyl-CoA atoms are colored by RMSF. Interestingly, the adenosine ring and the pantothenic acid groups of CoA are highly rigid inside the binding site. (B-D) RMSD time series of linoleoyl-CoA observed in every repetition of the MD simulations, presented by moiety: phospho-adenosine (B), pantothenate-cysteamine (C), and linoleoyl chain (D). (E) Histograms of distances between acyl-CoA and selected residues that contribute to binding. These results indicate that complementarity between the binding site and CoA is defined by a stacking of the adenosine ring with residue Trp205, hydrogen bonds with Ser303 and the backbone of Phe203 and electrostatic interactions between the 3' phosphate group of CoA and Arg204.

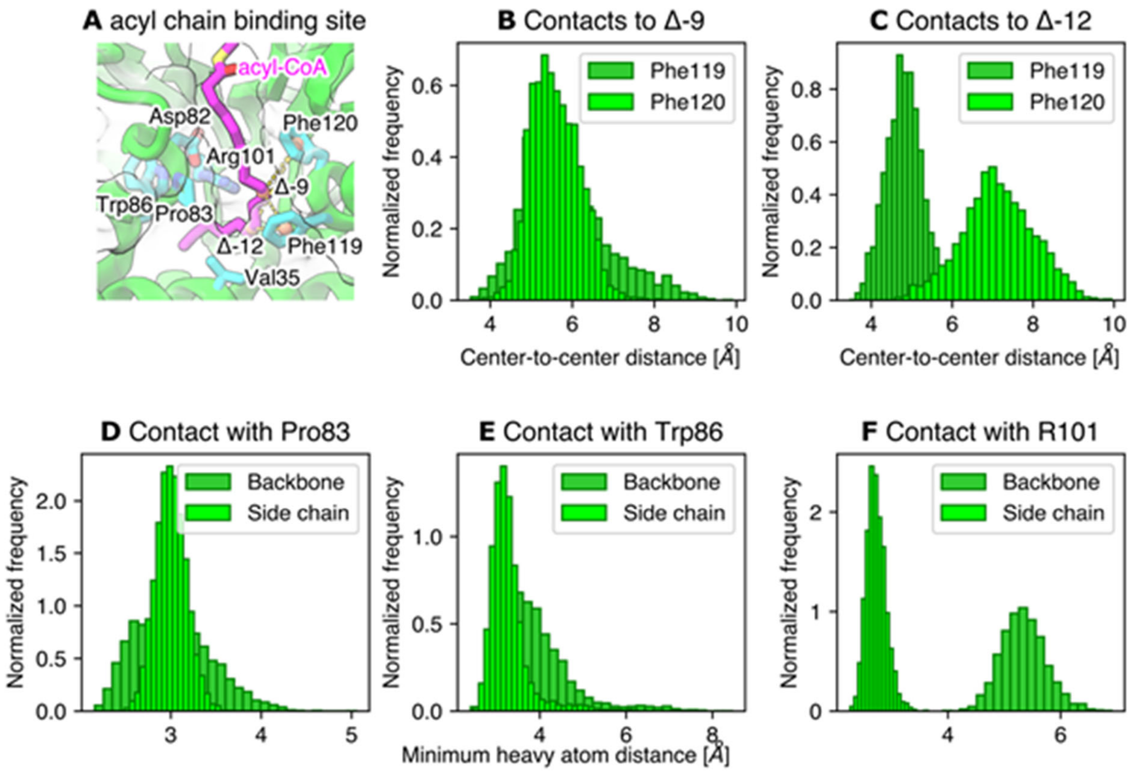


**Figure S6: Acyl chain binding to tafazzin.** (A) Conformation of the linoleoyl chain bound to the hydrophobic tunnel in tafazzin. (B-C) Histograms of the center of mass distances between each unsaturation in the bound linoleoyl chain and aromatic rings of residues Phe119 and Phe120. (D-F) Histograms of the minimum heavy atom distance between residues in the cavity and the activated acyl chain as sampled from simulations. The vertical, dashed lines correspond to ensemble averages.

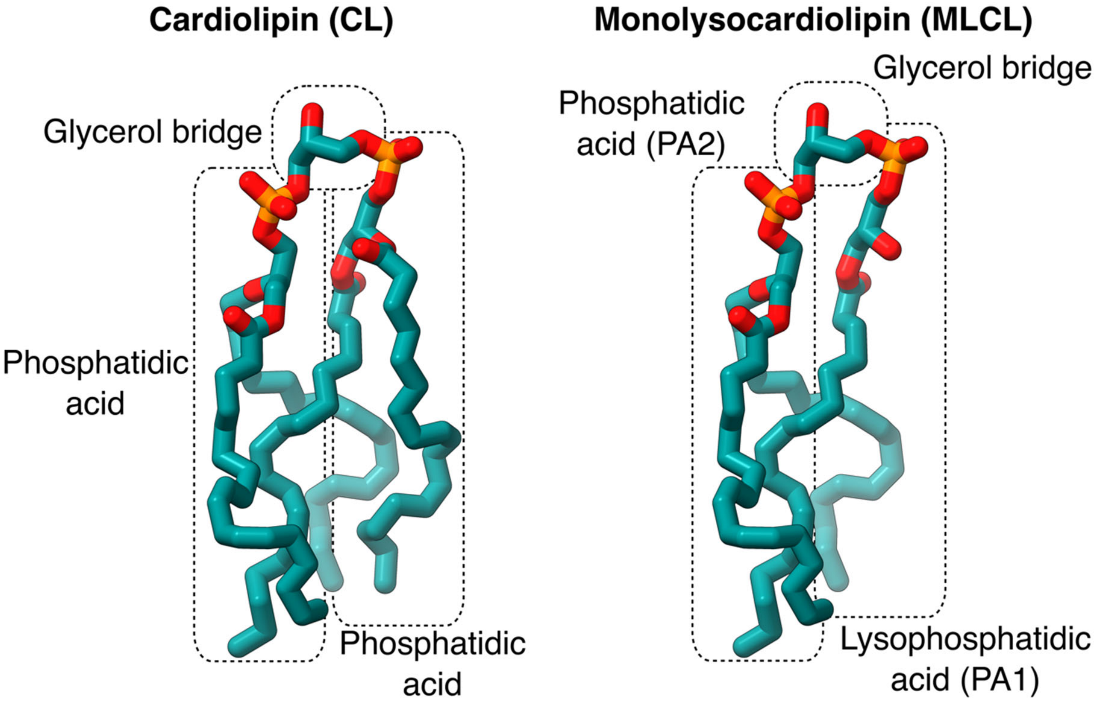


**Figure S7. General structure of cardiolipin and monolysocardiolipin and nomenclature used in this work.** Cardiolipin is formed by two phosphatidic acid subunits connected by a glycerol bridge. When all acyl chains are the same, both units are equivalent. In monolysocardiolipin, one of the units corresponds to a lysophosphatidic acid group (PA1) and the second unit is a phosphatidic acid molecule (PA2). PA1 contains the reactive hydroxyl group involved in the tafazzin-catalyzed reaction. Note that the free hydroxyl can be either in the *sn*-1 or *sn*-2 position.

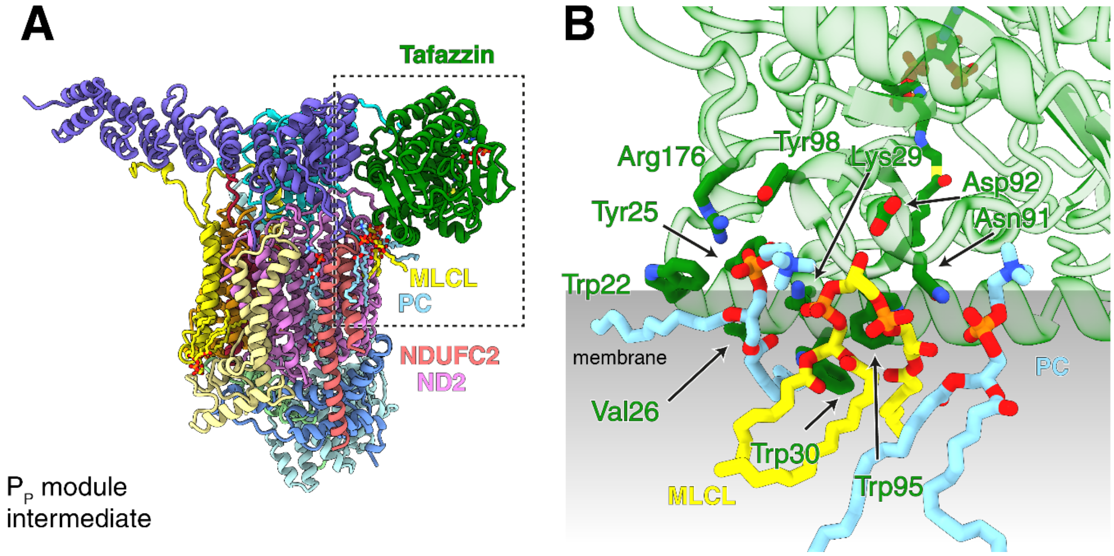


**Figure S8. Monolysocardiolipin bound to the interface of tafazzin/ND2/NDUFC2.** In the $P_P$ module assembly intermediate, monolysocardiolipin and two PC molecules are bound at the interface of tafazzin and the complex I subunits ND2 and NDUFC2. AlphaFold3 predictions match the structurally resolved lipids (compare Fig. 6). (A) position of the interface in the assembly intermediate. (B) Coordination of the lipids in a view similar to Figure 6.

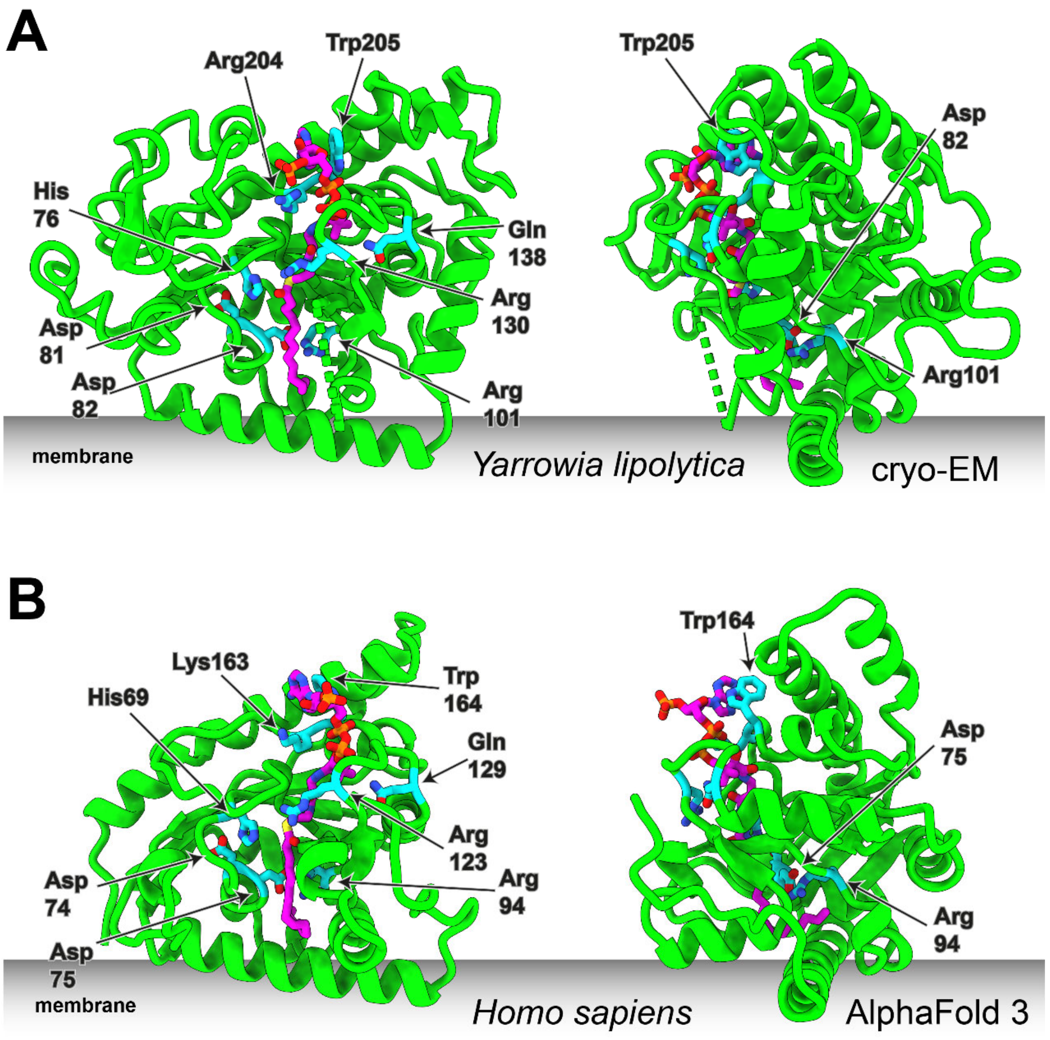


**Figure S9: Comparison of the *Y. lipolytica* cryoEM structure with the *H. sapiens* Alphafold 3 model.** Key structural features including binding of acyl-CoA are conserved. Selected residues shown in stick representation.

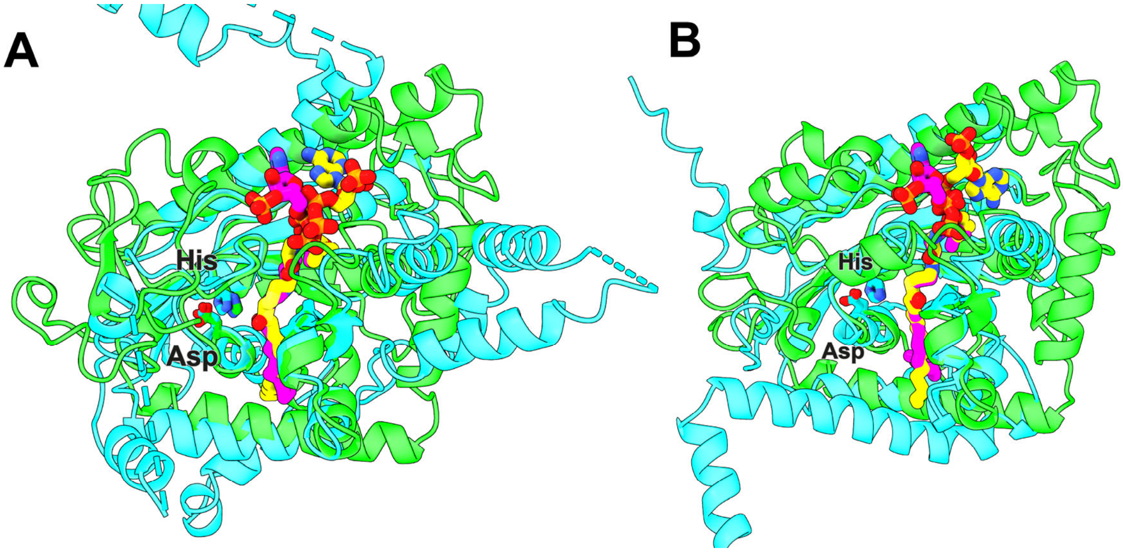


**Figure S10: Acyl-CoA binding sites of the AGPAT family.** Comparison of the binding positions of acyl-CoA in proteins of the AGPAT family. The conserved His$X_4$Asp motif is indicated for orientation. (A) Overlay of the two experimental structures of AGPAT proteins, GPAT1 (PDB:8E4Y; protein, cyan; 2-oxo-hexadecyl-CoA, yellow; only acyltransferase domain of GPAT1 shown) and tafazzin (cryo-EM structure described here, green; acyl-CoA, magenta). (B) Overlay of AlphaFold 3 models of LPLAT1 (cyan; acyl-CoA yellow) and tafazzin (green, acyl-CoA magenta). The phosphoadenosine head group of CoA is bound differently in tafazzin and the other two proteins, while binding positions for the central part of acyl-CoA match well.

**Tabel S1**: Barth syndrome missense mutations and corresponding residues in *S. cerevisiae* and *Y. lipolytica* [1,2]

| ***Homo sapiens*** | | ***Saccharomyces cerevesiae*** | ***Yarrowia lipolytica*** |
|---|---|---|---|
| residue numbering full sequence[3] | Barth syndrome missense muntantion (sequence missing exon 5)[4] | corresponding residues | corresponding residues |
| 17 | W17C | W24 | W24 |
| 28 | V28del, V28E | T35 | V35 |
| 29 | G29N | F36 | Y36 |
| 40 | N40D | Y47 | K46 |
| 43 | T43P | K50 | D49 |
| 50 | L50P | L57 | L56 |
| 54 | I54N, I54M | L61 | R61 |
| 57 | R57L | R63 | A63 |
| 62 | P62L | G70 | G69 |
| 69 | H69Q | H77Q | H76 |
| 71 | S71P | S79P | S78 |
| 74 | D74E | D82 | D81 |
| 75 | D75H, D75N | D83 | D82 |
| 76 | P76R | P84 | P83 |
| 77 | H77R | L85 | V84 |
| 80 | G80E or r.sp. | A88R/E | G87 |
| 82 | L82P | L90P | L89 |
| 94 | R94C, R94G, R94S, R94H, R94L | R102C/S | R101 |
| 101 | D101V, D101Y | N109V/D | D108 |
| 103 | C103R + path var MYH7 | C111 | C110 |
| 104 | F104V, F104L | F112V | Y111 |
| 110 | S110P | A118P | S117 |
| 116 | G116D | G124D | G123 |
| 117 | K117E | Q125E | Q124 |
| 118 | C118R | V126 | V125 |
| 119 | V119M, V119G | L127 | L126 |
| 120 | P120T | S128 | P127 |
| 124 | G124R | V134 | I133 |
| 159 | Q129P | Q138 | Q138 |
| 161 | G131R | S140R | G140 |
| 163 | D133G | D142 | D142 |
| 169 | L139F, L139H | L148H | L148 |
| 170 | W144G | W183 | W180 |
| 176 | H146Y,H146R, H146P | H185 | H182 |
| 178 | F148I, F148L | Y187I | F184 |
| 183 | V153G | V192G | V189 |
| 195 | G165V | G209 | G206 |
| 197 | G167R, G167W, G167VG167E, | T211 | G208 |
| 198 | R168L | R212 | R209 |
| 201 | A171V | L215 | L212 |
| 208 | I178V, I178N | I222 | I219 |
| 209 | I179N | V223D | I220 |
| 210 | L180R | V224R | V221 |
| 212 | L182P | I226P | M223 |
| 214 | H184R | A228 | S225 |
| 216 | G186R, G186E, G186V | G230R | G227 |
| 240 | G210R | G261 | G257 |
| 267 | T237K | A310 | A306 |

[1] red = disease associated (*H.s.)* or catalytically impaired variant (*S.c.*)
[2] yellow= active site, grey= acyl binding pocket, violet = near acyl chain cavity, purple= CoA headgroup
[3] full sequence including exon 5, Barth syndrome Database (https://www.barthsyndrome.org/)
[4] canonical splice variant of tafazzin

**Supplementary movies:** https://oc.biophys.mpg.de/owncloud/s/Qg6d37LpspQAafJ

**Movie S1**: **CoA coordination in Tafazzin**

CryoEM structure of the $P_P$ module intermediate of respiratory complex I with a focus on tafazzin and the residues coordinating acyl-CoA.

**Movie S2: MD simulation of acyl-CoA bound to tafazzin.**

The animation shows a sample trajectory with detailed information about the interactions between key residues in tafazzin responsible for recognition of acyl-CoA. To facilitate visualization, the binding site is divided following acyl-CoA fragments.

**Movie S3: MD simulation of tafazzin in complex with acyl-CoA and MLCL in explicit membrane.**

This movie shows the orientation of tafazzin in the inner mitochondrial membrane and describes the MLCL binding site predicted by AlphaFold 3. A sample trajectory from MD simulations is also shown to observe the binding and flexibility of MLCL in the protein.

**Movie S4: MD simulation of tafazzin in complex with CoA and PC in explicit membrane.**

The trajectory shows how PC occupies the same cavity where the acyl chain of acyl-CoA binds. In turn, CoA binds in a similar conformation to its acylated form. PC also occupies the A1 and A2 sites where MLCL is predicted to bind, indicating that there is no space for binding of a second phospholipid.

**Movie S5**: **Disease related residues**.

AlphaFold 3 model of tafazzin from *H. sapiens*, for mutations compare Table S1, details see text.
